# Unveiling Nucleosome Dynamics: A Comparative Study Using All-Atom and Coarse-Grained Simulations Enhanced by Principal Component Analysis

**DOI:** 10.1101/2024.11.05.622089

**Authors:** Abhik Ghosh Moulick, Rutika Patel, Augustine Onyema, Sharon M. Loverde

## Abstract

This study investigates nucleosome dynamics using both all-atom and coarse-grained (CG) molecular dynamics simulations, focusing on the SIRAH force field. Simulations are performed for two nucleosomal DNA sequences—ASP and Widom-601—over six microseconds at physiological salt concentrations. Comparative analysis of structural parameters, such as groove widths and base pair geometries, reveals good agreement between atomistic and CG models, though CG simulations exhibit broader conformational sampling and greater breathing motion of DNA ends. Principal component analysis (PCA) is applied to DNA structural parameters, revealing multiple free energy minima, especially in CG simulations. These findings highlight the potential of the SIRAH CG force field for studying large-scale nucleosome dynamics, offering insights into DNA repositioning and sequence-dependent behavior.

## 1. Introduction

Protein-DNA interactions are important for many essential processes inside the cell, such as organizing genetic information within the nucleus in eukaryotes, regulating gene expression, and DNA replication and transcription^1–3^. The nucleosome core particle (NCP), the elementary building block of chromatin, is a well-known protein-DNA complex that packages the genome inside eukaryotic cells^4–6^. The NCP is comprised of two copies of 4 histone subunits, i.e., H2A, H2B, H3, and H4. Each histone subunit consists of a highly ordered helical globular core region enclosed by flexible, disordered, positively charged tails known as the histone tails. The tails play an essential role in NCP-NCP interaction and the higher-level organization of chromatin. The NCP has a 147 base pair duplex DNA wrapped ∼ 1.7 times around a positively charged histone protein core in a left-handed helical fashion. The nucleosome represents 75-90% of the whole genome. It plays a pivotal role in several genomic processes, such as transcription, a process by which RNA polymerase copies DNA into RNA. During transcription, RNA polymerase must read the DNA sequence enclosed in the nucleosome. The timescale associated with transcription is from seconds to minutes. The positioning of nucleosomes along the DNA controls the accessibility to DNA binding factors, such as transcription factors and RNA polymerase^7^. Hence, understanding the underlying sequence dependence that governs nucleosome positioning and dynamics, particularly the breathing motion, is crucial to gaining a broader mechanistic insight into genomic processes.

Nucleosome dynamics are associated with an extended range of timescales, such as unwrapping nucleosomes that range from milliseconds to seconds. Small-scale rearrangements, such as breathing or loop formation, happen at microsecond time scales^8–12^. The repositioning of the DNA on the nucleosome surface is associated with timescales of minutes to hours^13–15^. Henceforth, a single experimental technique cannot cover the extended ranges of timescales. Atomistic molecular dynamics (MD) simulation using state-of-the-art forcefields can complement experimental findings, helping to decipher molecular details underlying these events. Atomistic MD simulation has already been employed in several studies based on the nucleosome, such as the role played by hydration patterns and counterions around the nucleosome^16^, the role of the histone tails^8, 17, 18^, and sequence-dependent nucleosome dynamics^19–24^. An increasing number of computational studies with atomistic details of nucleosomes are reported based on multiple microseconds time scales^25–27^. These studies have reported on the formation of twist defects^26^, loop formation^25^, etc. Characterizing higher-order nucleosome organization, such as the force between nucleosome dimers, the tetra nucleosome free energy landscape, or simualtions of chromatin fibers, requires implicit solvent approaches^21^ or coarse-grained models^28^. The plasticity of the nucleosome is also suggested to influence the phase behavior^29^. The underlying stability and structure of protein-DNA complexes largely depend on the accuracy of the forcefields^30^. Atomistic MD simulations of nucleic acid complexes typically utilize either AMBER^31–33^ or CHARMM^34, 35^ force fields. Improvement of nucleic acid (NA) force fields mainly focuses on refining glycosidic torsion and backbone parameters^31, 36, 37^ that may manifest deficiencies only over long simulation timescales. Both CHARMM and AMBER-based simulations of nucleic acids maintain the experimental double helical structure of DNA at tens of microseconds^38, 39^. However, some artifacts have been reported for simulations of longer dsDNA fragments with the CHARMM36 forcefield in terms of structural stability^40^. Conversely, AMBER-based simulation force fields show good agreement with experiments with some minor and reversible distortions^40^. Shaw and coworkers^41^ developed a new DNA force field, Des-Amber, with refined non-bonded parameters. However, this force field cannot capture the BI/BII state correctly, whose population plays a key role in the flexibility of DNA and its ability to bind with proteins. Further advancements in force field development and the integration of multiscale modeling approaches will be essential to overcome these limitations and accurately capture the full spectrum of nucleosome dynamics.

Molecular dynamics studies of the NCP performed in our laboratory have observed correlated DNA motion of the DNA ends^23^. Indeed, at physiological salt concentrations, timescales for nucleosome ‘breathing^42^’ are suggested to be at 0.1-1 ms. To accelerate the dynamics of DNA motion, we simulated the NCP at salt concentrations ∼ 10 times physiological ion concentrations with a 5 µs trajectory. We found DNA partial unwrapping starting with a spontaneous loop that forms in the SHL-5 region ^25^, similar in location to that reported by Bilokapic *et al*^43^. We further report on large-scale DNA motion for two different nucleosomal DNA sequences— the ‘Widom- 601’ and alpha satellite palindromic ‘ASP’ nucleosomal DNA sequences— based on 12 µs simulation trajectories on Anton 2 at a high salt concentration of 2.4 M^24^. The two sequences exhibit different pathways, with the ‘ASP’ sequence forming a loop, while the ‘Widom-601’ shows large-scale breathing motion. We find that the motion of the H2A and H2B tails plays a key role in loop formation, while the H3 tail plays a critical role in breathing. Post translational modifications (PTM) are also suggested to modify nucleosome breathing motion^44^. We further investigate the dynamics of the histone tails, considering the role of the acetylation of the histone tails^45^, characterizing how salt modulates their conformational dynamics. Chemically accurate coarse-grained models are necessary to probe the role of sequence in modulating nucleosome breathing at physiological salt concentration.

It is well known that a simplified or ‘a coarse-grained (CG)’ representation of a complex system like a protein-DNA complex is advantageous for characterizing the dynamics of these complexes and their phase behavior^29, 46^. Insight into various biological problems can be obtained by choosing a resolution that fits the length and timescale of interest^47^. Force-induced unwrapping of the nucleosome has been characterized via polymer bead-spring models^48^. CG simulation was further explored to characterize the tension-dependent free energy profile of DNA as a function of extension^49^. Sun et al.^50^ developed a CG model to characterize nucleosome phase separation with explicit divalent and polyvalent ions based on a ‘bottom-up’ coarse-grained model. The nucleosomal DNA is modeled as five beads representing every two base pairs; the histone protein is modeled as a single bead for each amino acid, and one bead represents one ion for all ion species. Chakraborty et al.^51^ developed a CG model known as COFFEE (Coarse-grained force field for energy estimation) based on a self-organized polymer model. This model was used to study the salt-induced unwrapping of the nucleosome. Apart from polymer-based models of the nucleosome, higher-resolution coarse-grained models have been introduced to study nucleosome dynamics. For example, the Schlick group^52, 53^ used Brownian dynamics (BD) simulation to simulate fibers with a mesoscale model of chromatin. In this model, the histone core is treated as a cylinder with 300- point charges distributed on its irregular surface^54^, linker DNAs is represented by 1 bead per 3 nm^55^, flexible histone tails^56^ are explicitly incorporated along with flexible linker histone^57^. Zhang et al.^22^ investigate nucleosome unwrapping by combining the associative memory, water-mediated, structure, and energy model (AWSEM) force field^58^ for protein and the 3SPN model^59^ for DNA. The sequence-dependence dynamics of the nucleosome were probed using CG modeling by de Pablo’s group^60^. They capture sequence dependence dynamics of the nucleosome and show that nucleosome repositioning occurs either by loop propagation or twist diffusion. Based on de Pablo’s CG model, Takada’s group shows further aspects of sequence-dependent repositioning dynamics^61–63^, demonstrating two sliding modes based on the nucleosomal DNA sequence^61^. Collepardo et al. have shown that DNA breathing can modify the nucleosome nucleosome interaction and promote liquid-liquid phase separation (LLPS)^29^.

Higher-resolution chemistry-based CG force fields like SIRAH and MARTINI have successfully described DNA-protein interactions^64, 65^. Parameterization follows one of the two main strategies: a bottom-up approach, where the model focuses on reproducing microscopic features based on a more theoretical model such as an atomistic or quantum mechanical model, or a top-down approach, where the model is built in such a way that it can reproduce a set of experimental macroscopic properties like surface tension and density^66, 67^. MARTINI uses a bottom-up strategy for bonded interactions and a top-down for non-bonded interactions as a parametrization strategy, while SIRAH uses a bottom-up structure-based approach. The limitation of the MARTINI model lies in base-pairing, which is not specific and requires an elastic network to keep dsDNA in its canonical representation^68^. However, the SIRAH CG DNA model does not require an elastic network. Furthermore, the model shows good agreement with the structural properties of DNA^69^. The SIRAH CG^70–72^ force field has also been applied to numerous biomolecular systems, including protein-nucleic acid complexes^73–77^. For example, Machado et al.^78^ use a hybrid CG atomistic approach to probe the conformational dynamics of the Lac repressor-DNA complex. Due to the versatility of the force field, modified parameters to include salt bridges, and previous success in characterizing protein-nucleic acid complexes, we choose the SIRAH force field to characterize the dynamics of nucleosomal DNA in the nucleosome core particle.

Understanding DNA dynamics on the base pair level gives crucial insight into the repositioning of DNA along the histone core. Here, we probe if nucleosome dynamics is sequence-dependent by comparing six-microsecond atomistic simulations with multiple replicas of the same systems using the SIRAH force field. We consider two different NCP nucleic acid sequences: (i) the human ⍺- satellite palindromic sequence (ASP) and (ii) the strong positioning ‘Widom-601’ DNA sequence. An earlier study using the SIRAH CG force field shows good agreement with atomistic simulation for the Drew-Dickerson dodecamer (DD) at the base pair level^79^. Motivated by this, we address base pair and local geometry, such as intra and inter-base pair parameters, for these two nucleosomal DNA sequences. First, we compare various structural parameters of the nucleosomal DNA based on the radius of gyration, groove width, and intra- and inter-base pair parameters. We find good structural similarity in atomistic and CG simulation base-pair parameters. Next, we quantify the breathing motion of DNA End-1 and End-2 for both atomistic and CG simulations. We find significant breathing motion at physiological salt concentration for CG simulations compared to AA simulations. We also characterize DNA repositioning around the histone protein in terms of translational and rotational order parameters, as first described by Lequieu et al.^80^ Overall, our study on the nucleosome core particle establishes the accuracy of the SIRAH CG force field in characterizing large-scale motion, including breathing of the DNA. We also demonstrate that this model can probe the translocation and rotation of the DNA in the nucleosome core particle. We demonstrate that methods in dimensionality reduction, such as principal component analysis (PCA), can be applied to DNA order parameters to extract conformations of the DNA where the breathing motion occurs, finding that these conformations correspond to key states in the translocation and rotational space of the free energy landscape.

## 2. Methods

### 2.1 System Preparation

Here, we consider two different sequences of nucleosome DNA in complex with the histone in the nucleosome core particle (NCP), (i) the human ⍺-satellite palindromic sequence (ASP) and (ii) the strong positioning ‘Widom-601’ DNA sequence. The initial coordinates for the ASP NCP are taken from the PDB ID of 1KX5^81^. The crystal structure of 1KX5 contains 14 Mn^2+^ ions. Because of the absence of force fields for Mn^2+^, we replace these ions with Mg^2+^. For the ‘Widom-601’ sequence, we consider the initial coordinates obtained from the protein data bank having PDB ID 3LZ0^82^. This crystal structure has missing histone N-terminal tails. So, we model these missing tails and other missing residues using Prime of the Schrodinger software suite, as previously reported^83, 84^. The ASP structure is used as the template for homology modeling. We replace the 8 Mn^2+^ ions in the crystal structure of the homology-modeled 3LZ0 system with Mg^2+^ ions to use the available force fields for Mg^2+^ ions.

### 2.2 All-atom Simulation of the NCP

Next, we simulate both NCP sequences using an all-atom molecular dynamics simulation using 0.15 M NaCl salt. The histone proteins are parametrized using the AMBER19SB force field^85^, whereas DNA is parametrized using OL15^86^. The OPC water model^87^ is used as solvent around the NCP in an orthorhombic box. Na^+^ and Cl^-^ ions are parametrized using Joung and Cheetham parameters (2008)^88^, while the Li/Merz compromised parameter set^89^ was used for Mg^2+^ ions. According to Kulkarni et al., the Lennard-Jones interaction of Na+/OPC (OW) was improved to better estimate osmotic pressure^90^. After parametrization, both systems are minimized for 15000 steps, following the steepest descent and conjugate gradients in the AMBER18 package^91^. Then, both systems are heated at constant volume, slowly varying the temperature to 310K. All bonds involving hydrogen atoms are constrained using the SHAKE algorithm^92^. The heated structures are further equilibrated for 100 nanoseconds (ns), maintaining a constant pressure of 1 Bar using a Berendsen barostat and a constant temperature around 310K using a Langevin thermostat with a collision frequency of 1.0 ps. The total electrostatic interaction is calculated using a Particle Mesh Ewald (PME) algorithm with full periodic boundary conditions. The cut-off value of 12 Å was considered for the van der Waals interaction, while bonded atoms were excluded from non-bonded atom interactions using a scaled 1-4 value. The Gaussian Split Ewald method was used to accelerate the electrostatic calculations. The final production runs are carried out for six µs on Anton 2^93^. The system-specific description is given in Table 1.

**Table 1.** Summary of initial set-up of both All-atom (AA) and Coarse-grained (CG) simulation.

| NCP systems | 1KX5-AA | 3LZ0-AA | 1KX5-CG | 3LZ0-CG |
| --- | --- | --- | --- | --- |
| Box dimensions, Å | 159 x 191 x 112 | 171 x 185 x 124 | 210 x 210 x 210 | 213 x 213 x 213 |
| No. of atoms | 444888 | 448776 | 114459 | 117837 |
| No. of solvent molecules | 104740 | 105816 | 26587 | 27448 |
| No. of Na<sup>+</sup> ions | 472 | 486 | 896 | 930 |
| No. of Cl<sup>-</sup> ions | 356 | 358 | 783 | 805 |
| No. of Mg<sup>2+</sup><br>ions | 14 | 8 | 14 | 8 |
| Salt<br>concentration | 0.15M | 0.15M | 0.15M | 0.15M |

### 2.3 Coarse-Grained Simulation of the NCP

Next, both NCP systems are simulated using the SIRAH coarse-grained forcefield in the GROMACS package^94^. Instead of the “four heavy atoms to one CG bead” rule according to the well-known CG MARTINI forcefield, the SIRAH CG force field handles the peptide bonds in the protein with a high level of detail by maintaining the coordinates of nitrogen (N), ⍺-carbon (C⍺), and oxygen (O). SIRAH models the side chain of the protein more coarsely. In the case of DNA, SIRAH reduces the complexity of nucleotides by considering six effective beads for each canonical nucleotide in DNA (A, T, C, and G). Each of the six nucleotide beads are placed in the exact cartesian coordinates of the corresponding atoms from the atomic representations. Two beads at the phosphate and C5’ carbon position represent the DNA backbone. The phosphate bead carries a-1 charge. Three beads represent the Watson-Crick edge. A-T and G-C base pairs identify each other through electrostatic complementarity. The partial charges add to zero on these CG beads at Watson-Crick edges. In the SIRAH CG representation, the details of Sugar moiety are completely ignored, with the 5-member ring replaced with one bead in the C1 position, which connects the backbone to the Watson-Crick edge. SIRAH uses a WT4 water model formed by four linked beads, each having a partial charge. This charge pattern is allowed to generate its dielectric permittivity. This CG water model can include ionic strength effects by including explicit salt and reproduces the osmotic pressure of water. To maintain the transferability between different MD packages, SIRAH uses the commonly found classical Hamiltonian function, which typically includes bonded (bond stretching, bending, torsion angle, etc.) and non-bonded (Lennard-Jones and Coulombic potentials).

Here, we simulate both NCPs following the protocol mentioned in Machado et al.^72^ for three sets of six-microsecond simulations for both nucleic acid sequences, the ASP and Widom-601. SIRAH tools were extensively used for mapping and analysis purposes. Before mapping to the CG model, the PDB2PQR server^95^ set the protonation state based on the assumption of neutral pH by the AMBER naming scheme. After mapping into the CG model, the protein-DNA complex is solvated in a cubic box of SIRAH WT4 water^96^. The system is neutralized by adding Na^+^ and Cl^-^ ion at 0.15 M salt concentration. The required number of ions, box dimensions, and total number of atoms and solvent molecules are tabulated in Table 1 for the 1KX5 and 3LZ0 structures. The box size is chosen to be sufficiently large so that the complex does not interact with its periodic image. Two steps of minimization are performed during system preparation. At first, the protein side chains are energy minimized by restraining the backbone for 50000 steps using the steepest decent algorithm. This step improves the structural stability of the protein by avoiding significant distortions to the secondary structure of the protein. Then, the whole system was energy minimized for 5000 steps following the steepest descent. Next, solvent molecules are equilibrated around the complex by simulating each complex for five ns while placing a harmonic restraint on the position of all CG beads. The temperature of the system is set at 310K using a V-rescale thermostat^97^.

To improve the solvation of protein side chains, a further 25 ns equilibration is performed, maintaining the temperature at 310K. Finally, unrestrained simulation is carried out for six µs maintaining pressure to 1 atm using Parrinello-Rahman Barostat with isotropic pressure coupling.

The time step for all the simulations is fixed at 20 fs. The Particle Mesh Ewald with a cut-off of 12 Å and a grid spacing of 2 Å is used for electrostatic interactions. For van der Waals interaction, the cut-off is set at 12 Å. All the parameters during the simulations are kept the same for the 1KX5 and 3LZ0 systems. Each system is simulated for three different replicas. All analyses are done by averaging all available replicas for each NCP system.

The back mapping from CG to All-atom is performed using the SIRAH Backmap^98^ tools. All the analyses are done on the obtained back-mapped trajectories to compare with all-atom trajectories. The atomistic positions in the back-mapped trajectory are built on a by-residue basis, maintaining the geometrical reconstruction (internal coordinates) following Parsons et al.^99^ The structures from the initial stage are protonated and minimized using the ff14SB^100^ atomistic force field within the tleap module of AmberTools^101^.

## 3. Analysis

Each analysis is performed for both NCP systems, comparing the all-atom and back-mapped trajectories obtained from coarse-grained simulation. In the rest of the text, “AA” denotes the all-atom trajectory, while “CG” is used for the back-mapped CG trajectory.

### 3.1 Radius of Gyration (R_g_)

We calculate the radius of gyration (R_g_) to compare the structures in the NCP in both AA and CG trajectory for both the protein and the DNA. We consider the backbone Phosphate (P) atom for DNA and the carbon C_⍺_ atom for the protein. R_g_ is defined as the average distance of P/ C_⍺_ atoms from their centers of mass (R_CM_). The square of R_g_ is defined as:

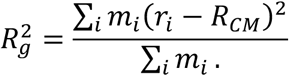

Here m_i_ and r_i_ is the mass and position of the i-th P/ C_⍺_ atom.

### 3.2 Secondary Structure Analysis

The secondary structure of the histone protein is analyzed for both CG and atomistic trajectories. For the atomistic trajectory, we used the AmberTools21^101^ secstruct tool, which employs the DSSP algorithm^102^. In DSSP, the hydrogen bonding pattern in the backbone amide (N-H) and carbonyl (C=O) positions determines the secondary structure of the protein. We use the sirah_ss tool of SIRAH tools to calculate the secondary structure for the CG trajectory. The secondary structure includes Helix, extended- β sheet and coil conformations. It calculates secondary structure based on hydrogen bond-like (HB) interactions and instantaneous values of the backbone’s torsional angles^71, 98^. The secondary structure propensity is calculated based on averaging over all trajectories for both CG and atomistic trajectories.

### 3.3 Structural properties for nucleosomal DNA

We evaluate well-known structural parameters applicable to DNA to compare AA and CG trajectories. These are (i) the major and minor groove width, (ii) the helical base pair step (inter base pair) parameters, and (iii) the helical base pair (intra base pair) parameters. All analyses were performed using the Curves+ software^103^. The inter-base pair parameters consist of three translations, i.e., shift (D_x_), slide (D_y_), and rise (D_z_), and three rotations, i.e., tilt (ϕ_x_), roll (ϕ_y_) and twist (ϕ_z_). Schematics are shown in Figure S1a. These parameters explain the relative position of two successive base pairs with respect to their short axis, long axis, and their normal.

We also calculate intra-base pair parameters, which comprise three translations, i.e., shear (S_X_), stretch (S_Y_), and stagger (S_Z_), and three rotations, i.e., buckle (θ_X_), propeller (θ_Y_), and opening (θ_Z_). Schematics are shown in Figure S1b. These parameters are calculated by determining the rigid-body transformations that map one base reference system to the others.

### 3.4 Principal Component Analysis

Principal Component Analysis (PCA) is a technique to characterize the collective motions of a molecule. It is a technique in dimensionality reduction by which one can identify configurational space having few degrees of freedom. This configurational space can be built by generating a 3Nx3N covariance matrix (C). Therefore, the C matrix is diagonalized where the elements of the matrix are represented as C = ⟨(q − ⟨q⟩)^T^(q − ⟨q⟩)⟩. Where q corresponds to coordinate and ⟨… ⟩ The bracket indicates the ensemble average. The diagonalization of this matrix gives i-th eigenvector and i-th eigenvalues. The projection of trajectory on the eigenvector provides the principal components (PC).

Here, we use dinucleotide base pair parameters as input coordinates for the PCA. The first two PCs were used to plot a two-dimensional free energy landscape. The free energy landscape can be obtained using the following equation: ΔG(PC1, PC2) = −k_B_Tln[P(PC1, PC2)/P_max_]. Here, Δ*G* represents the free energy of the state. P(PC1, PC2) is the joint probability distribution for PC1 and PC2, while k_B_ is Boltzmann’s constant and T is the temperature. P_max_ represents the maximum probability density.

### 3.5 Nucleosome Dynamics

#### 3.5.1 Breathing Motion of Nucleosomal DNA

We characterize the breathing motion of the nucleosomal DNA occurring in the DNA end regions due to the transient opening/closing of DNA entry/exit regions or in between the inner gyres, where two gyres come closer or move away from each other due to the modulation of histone-DNA contacts. We quantify the scope of DNA end breathing by calculating the breathing distance in the simulated structure, defined as the distance between the center of mass of SHL0 bp and the terminal bp present in the entry/exit region. Here, we represent the change in end breathing w.r.t the crystal structure. Positive values of end breathing distance indicate outward breathing w.r.t crystal structure, while negative values indicate inward breathing. We further quantify the breathing motion ^24, 25^ by calculating the displacement of each bp’s average distance (ΔR) over the last 3 µs of the simulated trajectories relative to the center of mass of nucleosomal DNA non-hydrogen atoms in the crystal structure.

#### 3.5.2 Translocation and Rotational Order Parameter

To quantify the movement of nucleosomal DNA around the histone protein, we observe the translocation and rotation of DNA position relative to the protein dyad through the translocation order parameter (S_T_) and the rotational order parameter (S_R_).^80^ Here, S_T_ is defined as,

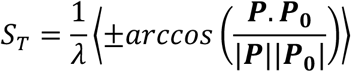

Here, **P** is a vector for a specific base pair which connects the histone center of mass to the center of mass of the respective base pair. **P_0_** is the value of the respective **P** in the crystal structure. λ is a conversion factor that converts radians into the base pairs of DNA translocation. The value of λ is 0.08rad/bp, mentioned in Ref.^80^ The sign of S_T_ is positive if (**P** x **P_0_)**.**f ≤** 0 (negative if > 0),

where **f** is a vector whose direction is along the center of the nucleosomal DNA superhelix. The positive value of S_T_ signifies forward translocation of nucleosomal DNA towards the 5’ end, whereas the negative value describes backward translocation towards the 3’ end. The schematic is shown in Figure S1c.

The S_R_ order parameter due to the rotational position of DNA is defined as,

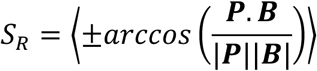

Here, **B** is a vector connecting the center of the given base step on the sense strand to its complementary base step on the antisense strand. All other terms are defined the same way S_T_. The value of S_R_ is positive if (**P** x **B)**.**D ≤** 0 (negative if > 0), where **D** is a vector from the 5’ to 3’ direction along the sense strand. If S_R_ = 1/2, then the minor groove is oriented away from the histone core, whereas SR = -1/2 signifies the orientation of the minor groove towards the histone core. The schematic is shown in Figure S1d. S_T_ and S_R_ order parameters have been used earlier to quantify the spatial positioning of DNA around histone proteins^80^.

#### 3.5.3 Minimum free energy path calculation

To identify the minimum free energy path between two conformations over a 2D free energy surface, we use the string method^104^ as implemented in MEPplot^105^. This method describes the pathway between two conformational states as a discrete set of points (known as beads) that evolve iteratively until they converge to a minimum free energy path. First, we identify two initial conformations from two different energy minima of 2D free energy surface. Finally, we obtain a path between two conformations using a gradient descent method where each point moves in the direction of the local gradient of the free energy surface in an iterative way.

## 4. Results

We perform comparative simulations of two well-known sequences of the NCP with the SIRAH force field and compare them against fully atomistic simulations. We use the ASP and the Widom-601 NCP sequences. Fig. 1a shows the ASP sequence’s crystal structure and its coarse-grained representation. The orientation of the nucleosomal DNA base pairs is represented with respect to the central base pair, commonly known as superhelical location (SHL) zero. In general, each SHL contains approximately 10 base pairs. It starts with SHL0 and ends at SHL ±7. Fig.1b shows the comparison of the sequence in nucleosomal DNA for both the ASP and the Widom-601 sequences. Several flexible dinucleotide steps, such as TA in the minor groove block, exist for the Widom-601 sequence, forming narrow conformations of the DNA. Both the minor grooves at SHL ±1.5 for the Widom-601 sequence contain the strong positioning motif TTTAA, which enhances its positioning affinity. Overall, there is a 15% greater G|C content in the Widom-601 sequence than in the ASP sequence. However, both sequences have similar G|C content in the minor grooves. Notably, the G|C content in the 601-R and 601-L halves of the DNA are different, with the right half containing a higher G|C content, which is thought to make it more rigid with fewer contacts with the DNA, and easier to open up under force as shown by Ngo et al^12^. Overall, the presence of G|C content and the strong positioning motif TTTAA in both SHL ±1.5 makes the Widom-601 one of the strongest positioning nucleosome sequences.

**Figure 1.**
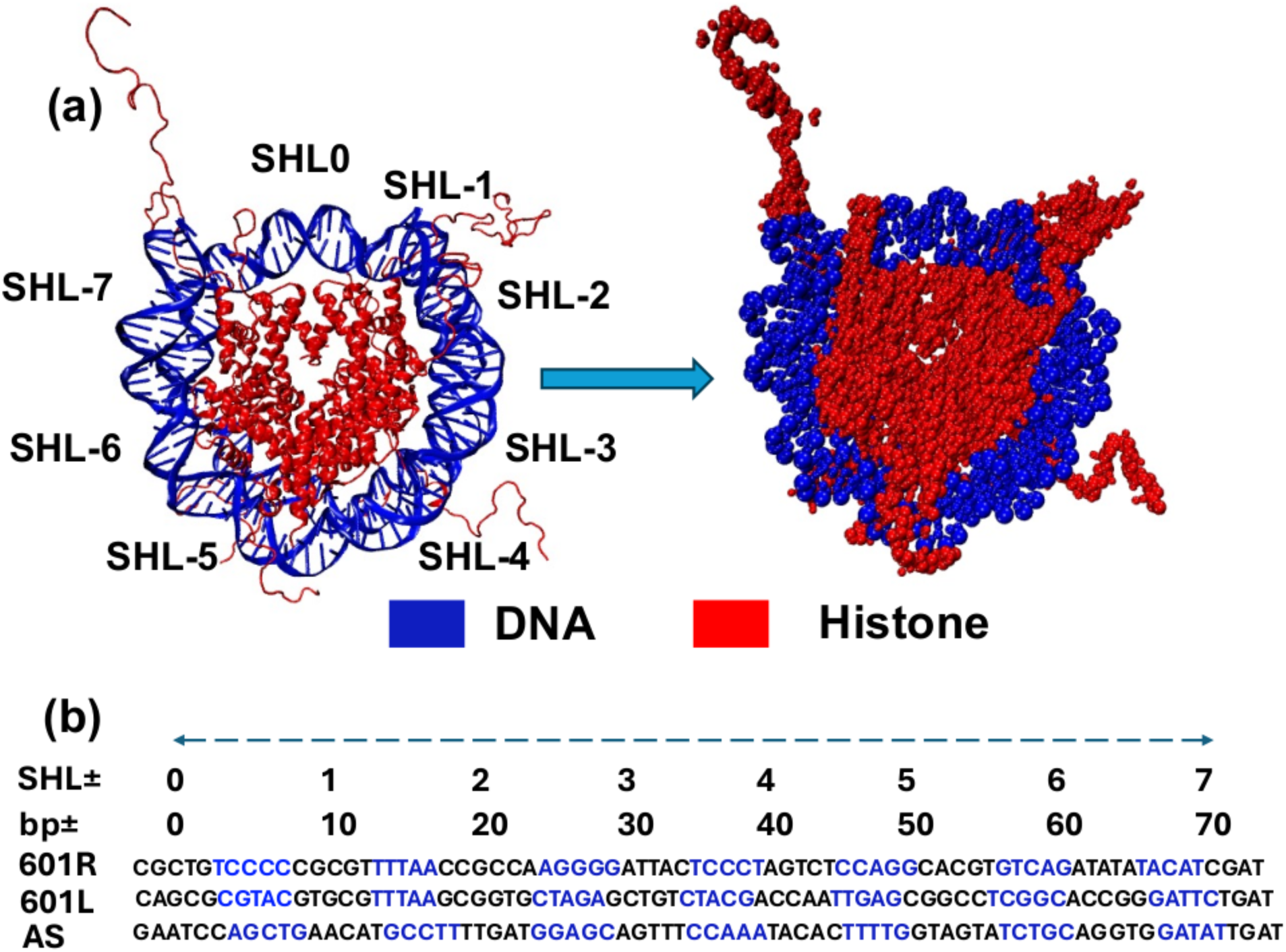
(a) Crystal structure of human ⍺-satellite palindromic sequence (ASP) sequence (PDB ID:1KX5) and its coarse-grained representation. DNA is marked in blue, while histone is marked in red. (b) Comparison of DNA sequence for Widom-601 and human ⍺-satellite sequence (ASP). The blue indicates a minor groove in the DNA sequence, while the black represents a major groove. Both halves of the Widom-601 (601-R and 601-L) sequence are shown, while for the ASP sequence, only one half is present as it is a palindromic sequence.

### 4.1 Structural comparison

A comparison of the radii of gyration (R_g_) of both fully atomistic and CG trajectories shows a direct comparison of the DNA at coarse-grained and all-atom levels. Fig. 2a shows the change of R_g_ over time for the ASP DNA sequence. Here, we characterize three independent replicas of CG trajectories (Rep1, Rep2, Rep3) with a single trajectory using all-atom force fields (AA). In Fig. 2b, we present the R_g_ histogram for all-atom and coarse-grain trajectories. The blue line indicates the average histogram over three independent CG trajectories. The average values of the R_g_ for the DNA over AA and CG trajectories are 45.69 ± 0.06 Å and 47.37 ± 0.05 Å, respectively. Fig. 2c depicts the overlapped equilibrium conformation of DNA for both the AA (green) and the CG (blue) trajectories. The values of R_g_ for the equilibrium AA and CG structures are 45.77 Å and 46.13 Å, respectively. We further compare the R_g_ of the DNA over time for the Widom-601 sequence (Fig. 2d). Fig. 2e illustrates the distribution of R_g_ for that sequence. The average R_g_ values are 45.52 ± 0.03 Å for CG and 47.39 ± 0.05 Å for AA. A representative equilibrium conformation for the Widom-601 DNA sequence is shown in Fig. 2f. Here, the values of R_g_ for AA and CG structures are 45.64 Å and 46.08 Å, respectively. The average R_g_ value of DNA obtained using the CG SIRAH force field for both sequences increases compared with the AA force field, indicating that the DNA samples have more conformational states in the coarse-grained trajectories.

**Figure 2.**
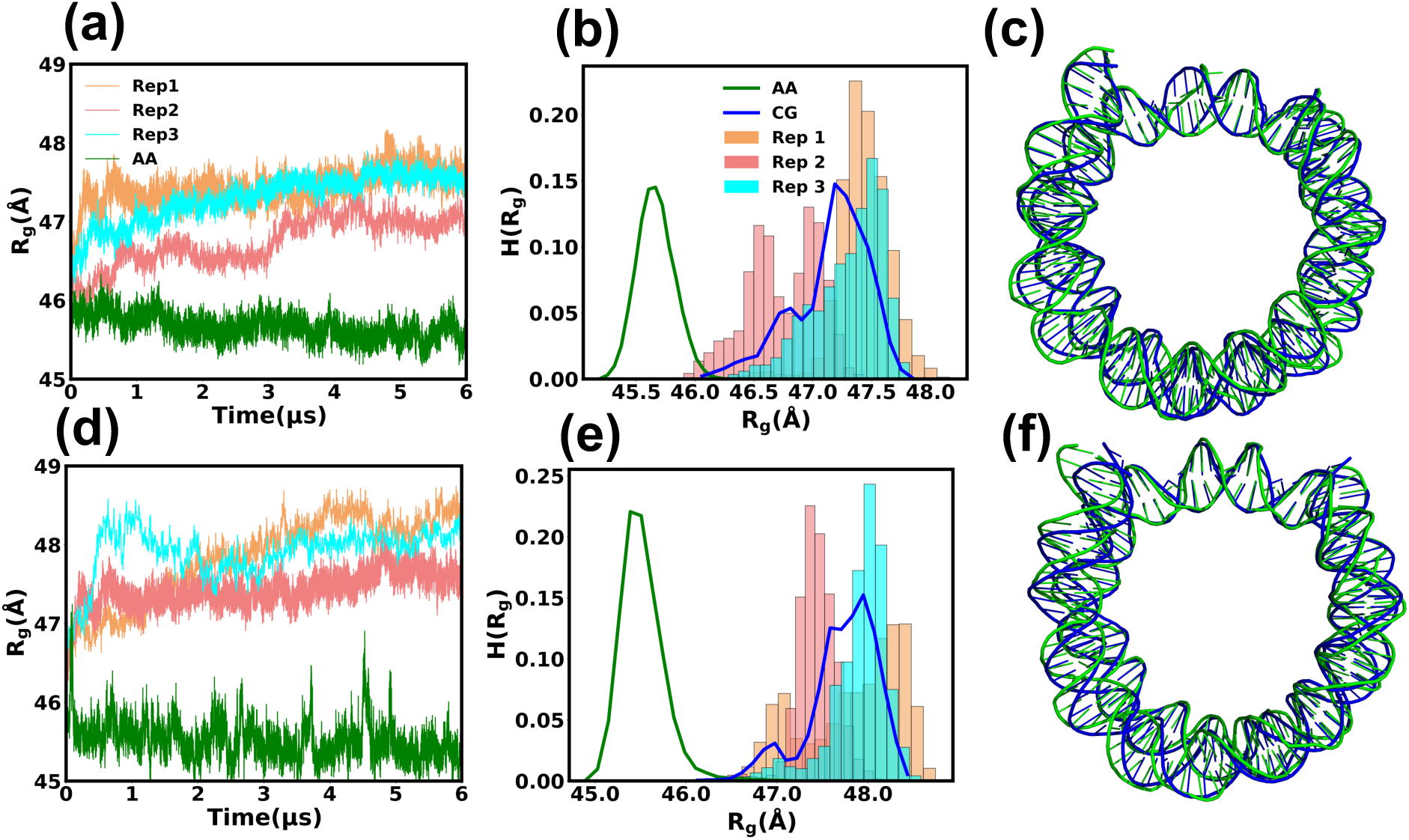
(a) Time evolution of radius of gyration (R_g_) of DNA considering phosphate atom for the ASP sequence. Results for three different replicas for coarse-grained trajectory and all atom trajectory are shown. (b) Histogram of R_g_ for different replicas and atomistic data. (c) Representative structure for 1KX5 DNA. (d) Time evolution of R_g_ of DNA for the Widom-601 sequence. Both CG replicas and atomistic simulation data is present. (e) Histogram of R_g_ for the Widom-601 sequence. (f) Representative overlapped structure for the Widom-601 DNA. Blue represents the backmapped atomic structure of CG trajectory while green represents the structure obtained using atomistic simulations.

Next, we compare the R_g_ of the histone protein, considering the C_⍺_ atom at different levels of detail. Fig. S2a illustrates the change in the R_g_ over time for the ASP histone protein, displaying three independent replicas of CG trajectories alongside trajectories using the all-atom force fields. In Fig. S2b, we present the histograms of R_g_ for both trajectory types. The average R_g_ values for the histone over the AA and CG trajectories are 34.26 ± 0.03 Å and 37.09 ± 0.14 Å, respectively. Fig. S2c depicts the overlapped equilibrium conformation of the histone for both the AA (green) and CG (blue) trajectories. We compare the R_g_ of the histone over time for another NCP sequence, the Widom-601, in Fig. S2d. Fig. S2e further details the distribution of R_g_ for that sequence, with average R_g_ values of 34.04 ± 0.1 Å for AA and 36.54 ± 0.09 Å for CG. S2f shows an overlapped equilibrium conformation of the histone proteins. For the histone, the average R_g_ value based on the CG force field shows good agreement with the atomistic force field results. Although similar to the DNA, the distribution of R_g_ states sampled for the protein CG trajectories is broader than the AA counterparts. Next, we compare the secondary structure percentage over the CG and AA trajectories. Fig.S2g shows the average percentage of the helix, extended, and coil conformation for the ASP sequence. The percentage of helix conformation is lower for the CG compared to the AA simulation. The extended and coil conformation percentage is higher for the CG simulation than for the atomistic counterpart. A similar scenario also holds for the Widom-601 (Fig. S2.h), i.e., the lower helical percentage in CG and a higher percentage of extended and coil conformations compared to atomistic simulation.

Next, we characterize the DNA structure regarding groove width and dinucleotide base-pair step parameters. Fig. S3a shows the schematic of DNA major and minor groove width over the overlapped equilibrated DNA conformation for both AA (green) and CG (blue) trajectories. The distribution of major groove width (d_Majw_) for the ASP DNA (Fig. S3b) suggests larger widths for the CG trajectories (blue) with an average value of 11.88 ± 0.07 Å as compared to the AA trajectory (green). The average d_Majw_ over the AA trajectory is 11.44 ± 0.04 Å. The average minor groove width (d_Minw_) for the ASP DNA over the CG trajectory and the AA trajectory is 5.44 ± 0.02 Å and 5.8 ± 0.01 Å, respectively. Fig. S3c depicts that the distribution peak of minor groove width distribution is lower for the CG (blue) than the AA (green) trajectory. The distribution of groove widths for the Widom-601 shows similar behavior as the ASP sequence for d_Majw_ (Fig. S3d) and d_Minw_ (Fig. S3e). The average d_Minw_ is 5.63 ± 0.01 Å over the CG trajectory, while for the all-atom trajectory, the average d_Minw_ is 5.68 ± 0.02 Å. Along the CG trajectory, the d_Majw_ average is 11.67 ± 0.01 Å, slightly higher than the average of 11.43 ± 0.01 Å observed over the all-atom trajectory. The similarity in major and minor groove widths suggests that the SIRAH coarse-grain force field can reliably approximate the groove widths of the DNA in both systems.

Next, to better understand the orientation of the DNA at the base pair level, we focus on the DNA inter-base pair parameters, which provide valuable insight into the structure and function of DNA molecules. Fig. 3 shows a histogram of different inter-base pair parameters obtained from CG and AA trajectories for the ASP DNA sequence. Table 2 tabulates the average inter-base pair parameter values obtained from CG and AA trajectories. The distributions of shift (D_X_) (Fig. 3a) parameters obtained from CG (blue) and AA (green) trajectories show close overlap. The average D_X_ value obtained from CG trajectories is 0.02 Å, whereas for AA trajectories, it is 0.0006 Å (See Table 2). Conversely, while the distributions of slide (D_Y_) (Fig. 3b) parameters and rise (D_Z_) parameters (Fig. 3c) from CG and AA trajectories did not overlap, the average value of these parameters across CG and AA trajectories show minimal disparity (See Table 2). Fig. 3d-f shows a histogram of rotational inter-base pair parameters, i.e., tilt (ϕ_X_) (Fig. 3d), roll (ϕ_Y_) (Fig. 3e), and twist (ϕ_Z_) (Fig. 3f). The histogram of tilt for CG (blue) and AA (green) trajectories exhibits complete overlap. CG trajectories yield an average tilt value of -0.28°, whereas the AA trajectory stood at -0.14° (see Table 2). The average twist value over the CG and AA trajectory is 32.00° and 34.02°, respectively. The roll order parameter shows distinct behavior as compared to the other parameters. The average value of roll over the CG trajectory is -8.42°, while for the AA, the average value is 2.19°. Fig.4 shows similar distributions of inter-base pair parameters for the Widom-601 sequence. The distribution of shift (D_X_) parameter (Fig. 4a) for CG overlaps with the AA trajectory. The average shift value obtained from CG is nearly equal to the AA average (Table 2). While the distributions of the slide (Fig. 4b) and rise (Fig. 4c) parameters from CG and AA trajectories do not overlap, the average values of these parameters along CG and AA trajectories show minor deviations. The rotational inter-base pair parameter tilt exhibits perfect overlap in distributions between the CG and AA trajectories (Fig. 4d). Additionally, the average value of the twist parameter (Fig. 4f) over CG and AA trajectories is 31.62° and 34.33°. The roll parameter shows similar behaviours as the ASP sequence. The distributions of roll over the CG and the AA simulations are shown in Fig. 4e. The average roll over the CG trajectories is -7.64°, while for the AA trajectory, it is 1.57°. Generally, the agreement of inter-base pair parameters between the CG and AA force fields is good. The deviation is mainly observed for the roll order parameter for both sequences.

**Figure 3.**
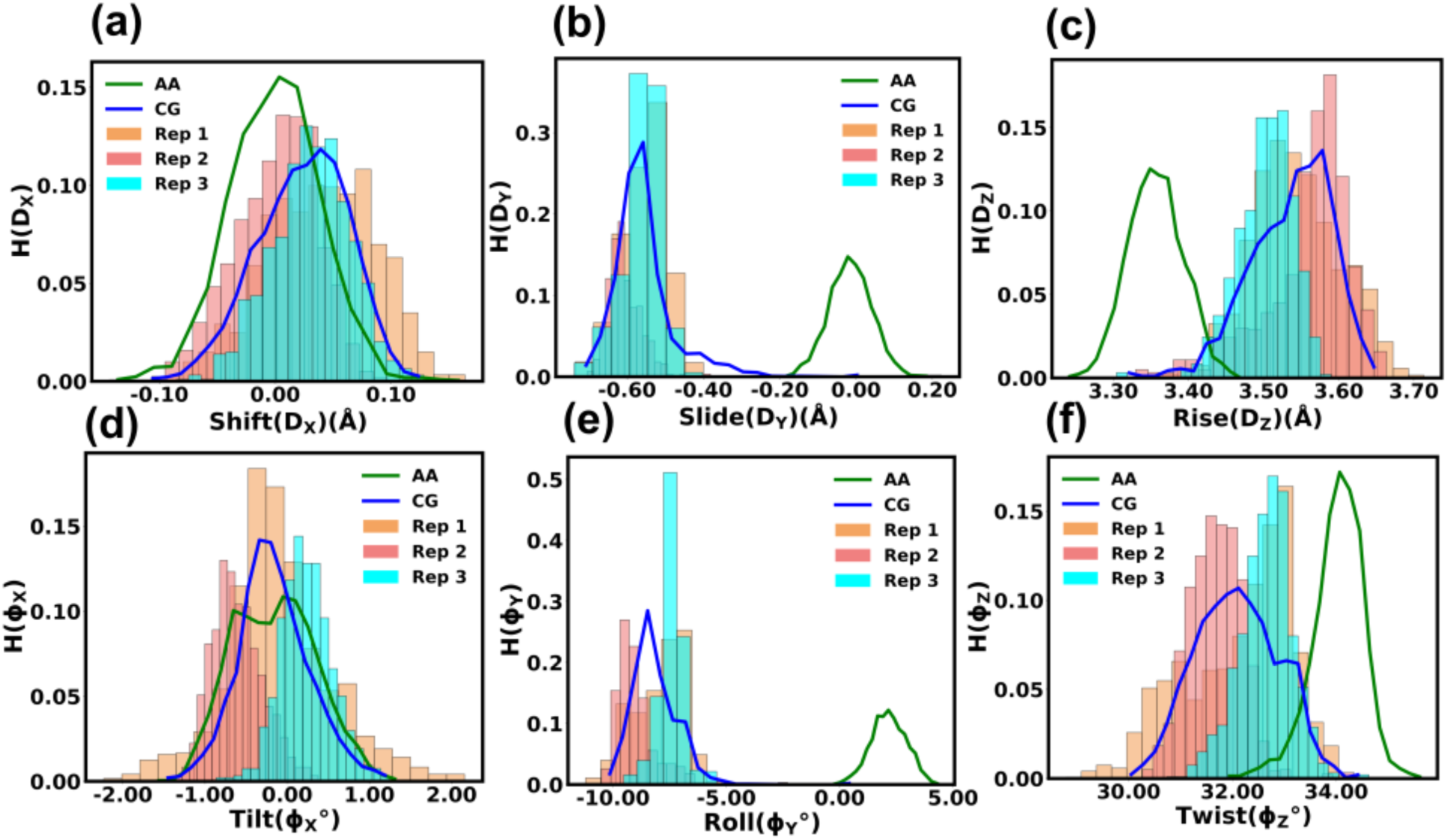
The histogram of DNA inter-base pair parameters for the ASP sequence: (a) Shift, (b) Slide, (c) Rise, (d) Tilt, (e) Roll, (f) Twist. Results for both atomistic and three different CG replicas are shown. Mean and errors are tabulated in Table 2.

**Figure 4.**
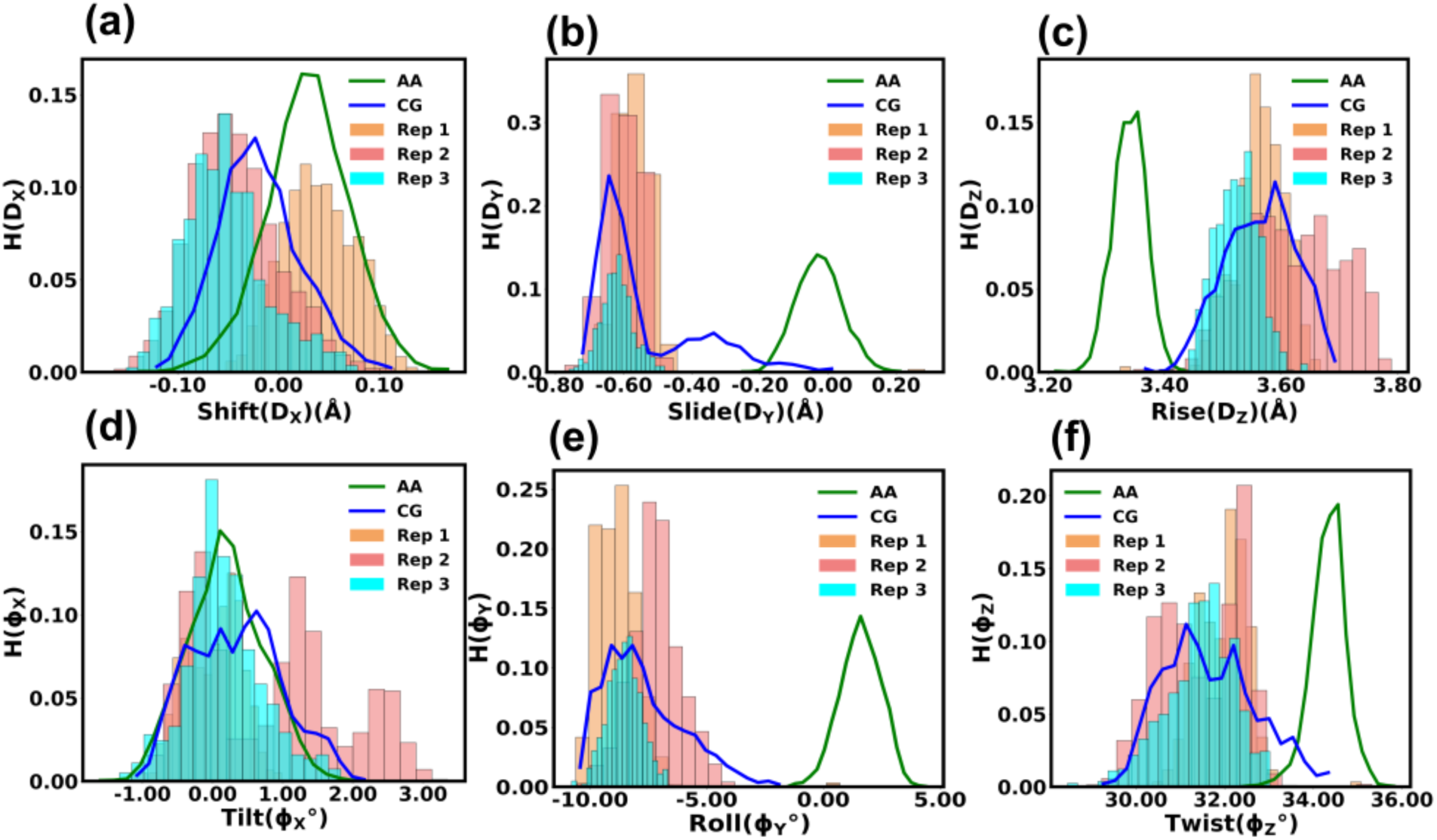
The histogram of DNA inter-base pair parameters for the Widom-601 sequence: (a) Shift, (b) Slide, (c) Rise, (d) Tilt, (e) Roll, (f) Twist. Results for both atomistic and three different CG replicas are shown. Mean and errors are tabulated in Table 2.

**Table 2.** Average values of inter-base pair parameters obtained from AA and CG trajectories. Error value is shown in parentheses.

| **NCP systems** | **1KX5-CG** | **1KX5-AA** | **3LZ0-CG** | **3LZ0-AA** |
| --- | --- | --- | --- | --- |
| Shift (D<sub>X</sub>) (Å) | 0.02 (0.01) | 0.0006 (0.001) | -0.02 (0.01) | 0.03 (0.003) |
| Slide (D<sub>Y</sub>) (Å) | -0.58 (0.01) | -0.01 (0.008) | -0.59 (0.02) | -0.03 (0.02) |
| Rise (D<sub>Z</sub>) (Å) | 3.54 (0.01) | 3.36 (0.02) | 3.6 (0.02) | 3.34 (0.004) |
| Tilt (φ<sub>X</sub>°) | -0.28 (0.14) | -0.14 (0.05) | 0.55 (0.14) | 0.19 (0.02) |
| Roll (φ<sub>Y</sub>°) | -8.42 (0.3) | 2.19 (0.11) | -7.64 (0.37) | 1.57 (0.08) |
| Twist (φ<sub>Z</sub>°) | 32.00 (0.36) | 34.02 (0.11) | 31.62 (0.19) | 34.33 (0.03) |

Next, the structural comparison between CG and AA is examined based on intra-base pair step parameters. Table 3 tabulates the average values of the intra-base pair parameters for the CG and AA trajectories. Fig. S4a shows distributions of the shear parameters for CG (blue) and AA (green) trajectories for the ASP sequence. The average value of the parameter over the trajectory for CG is 0.13 Å, while for AA, the value is 0.03 Å (Table 3). The conformations sampled for the CG trajectory are much broader than those for the AA trajectory. The distribution overlaps for the stretch parameter (Fig. S4b), although the CG trajectory exhibits a significantly broader range of conformations than AA. In the CG trajectory, the parameter averages -0.02 Å, while for the AA trajectory, the average value is 0.03 Å. The distribution of the stagger parameter (Fig. S4c) for CG and AA does not overlap, although the average value of stagger over CG and AA trajectory is 1.50° and 0.02°, respectively. The rotational intra-base pair parameter buckle exhibits overlaps between the AA (green) and the CG (blue) trajectory (Fig. S4d). The propel parameter shows distinct behaviors for the AA and CG simulations (Fig. S4e). The average value of the propel parameter for CG is -2.57°, while for AA, the value is -13.05°. For the opening parameter (Fig. S4f), the average value for CG is 8.00° and for AA is 2.85°. Most rotational inter-base pair parameters show good agreement in the average value along CG and AA trajectories, except propel and opening. We further investigate the inter-base pair step parameter for the Widdom-601 sequence. Fig. S5 shows the distribution of the parameters for both the CG and the AA trajectories. The distributions of the CG and AA trajectories partially overlap for shear (Fig. S5a) and stretch (Fig. S5b). The average value of both quantities along the CG and AA trajectories is nearly equal (Table 3). The distribution (Fig. S5c) does not overlap for the stagger parameter, although the average value for the CG is 0.98 Å and for the AA is 0.07 Å. The distribution for the buckle parameter (Fig. S5d) overlaps for CG and AA. The average propel parameter (Fig. S5e) value for CG is -0.56°, contrasting with AA’s -11.68°. As for the opening parameter (as depicted in Fig. S5f), CG averages 5.59°, whereas AA averages 2.32°. Most intra-base pair parameters exhibit consistent average values along the CG and AA trajectories, except for propel and opening, where notable differences are observed, like the ASP sequence.

**Table 3.** Average values of intra-base pair parameters obtained from AA and CG trajectories. Error value is shown in parentheses.

| **NCP systems** | **1KX5-CG** | **1KX5-AA** | **3LZ0-CG** | **3LZ0-AA** |
| --- | --- | --- | --- | --- |
| Shear ($S_X$) (Å) | 0.13 (0.01) | 0.03 (0.01) | -0.12 (0.01) | -0.02 (0.01) |
| Stretch ($S_Y$) (Å) | -0.02 (0.05) | 0.03 (0.006) | 0.01 (0.02) | 0.07 (0.03) |
| Stagger ($S_Z$) (Å) | 1.50 (0.008) | 0.02 (0.02) | 0.98 (0.02) | 0.07 (0.009) |
| Buckle ($\theta_X^\circ$) | 0.95 (0.44) | -0.49 (0.08) | 0.63 (0.26) | 0.77 (0.06) |
| Propel ($\theta_Y^\circ$) | -2.57 (0.16) | -13.05 (0.16) | -0.56 (0.6) | -11.68 (0.17) |
| Opening ($\theta_z^\circ$) | 8.00 (0.88) | 2.85 (0.07) | 5.59 (0.54) | 2.32 (0.07) |

### 4.2 Breathing Motion of Nucleosomal DNA

Here, we quantify the extent of End-breathing by computing the breathing distance in the simulated structure, defined as the distance between the center of mass of SHL0 bp and the terminal bp present at the entry/exit region. We compare the breathing motion of the nucleosomal DNA ends for both sequences. Fig. 5 shows a histogram of the breathing distance for both End1 and End2, depicted as the difference with respect to the crystal structure. For the atomistic trajectory, the breathing distance for both the DNA ends (Fig. 5a-b, marked in green) fluctuates near zero for the ASP sequences. The average value of the breathing distance for End1 is 1.22 Å, and for End2, it is 0.46 Å. For the CG trajectory, the breathing distance increases for both Ends (Fig. 5a-b, marked in blue). The average breathing distance for End1 is 17.69 Å, and for End2, it is 6.31 Å. The extent of breathing for both ends is different, with End1 displaying more extensive breathing since breathing motion is asymmetric, as suggested by earlier theoretical and experimental studies^11, 22,23^. Fig. 6 furhter displays the time evolution of breathing distance for both End-1 and End-2, represented as the difference with respect to the crystal structure. Fig.6 a-c shows snapshots of the nucleosomal DNA at different times from the atomistic simulation, indicating negligible breathing motion.

**Figure 5.**
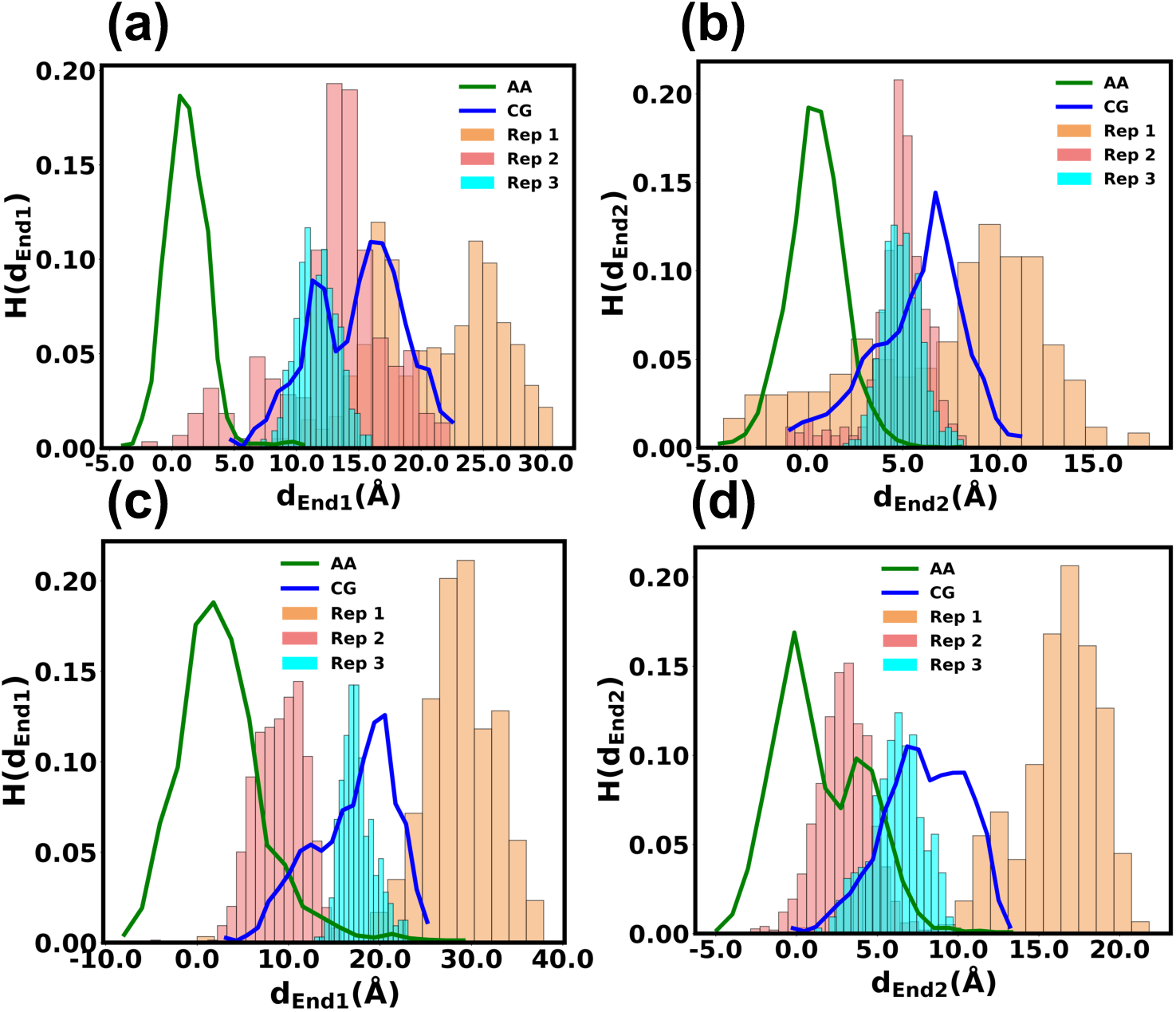
Normalized probability distribution of the breathing distance for the nucleosomal DNA for (a) End1 (ASP-L), (b) End2 (ASP-R) for the ASP sequence and (c) End1 (601-L), (d) End2 (601-R) for the Widom-601 sequence. Results for both the atomistic and three different CG replicas are shown.

**Figure 6.**
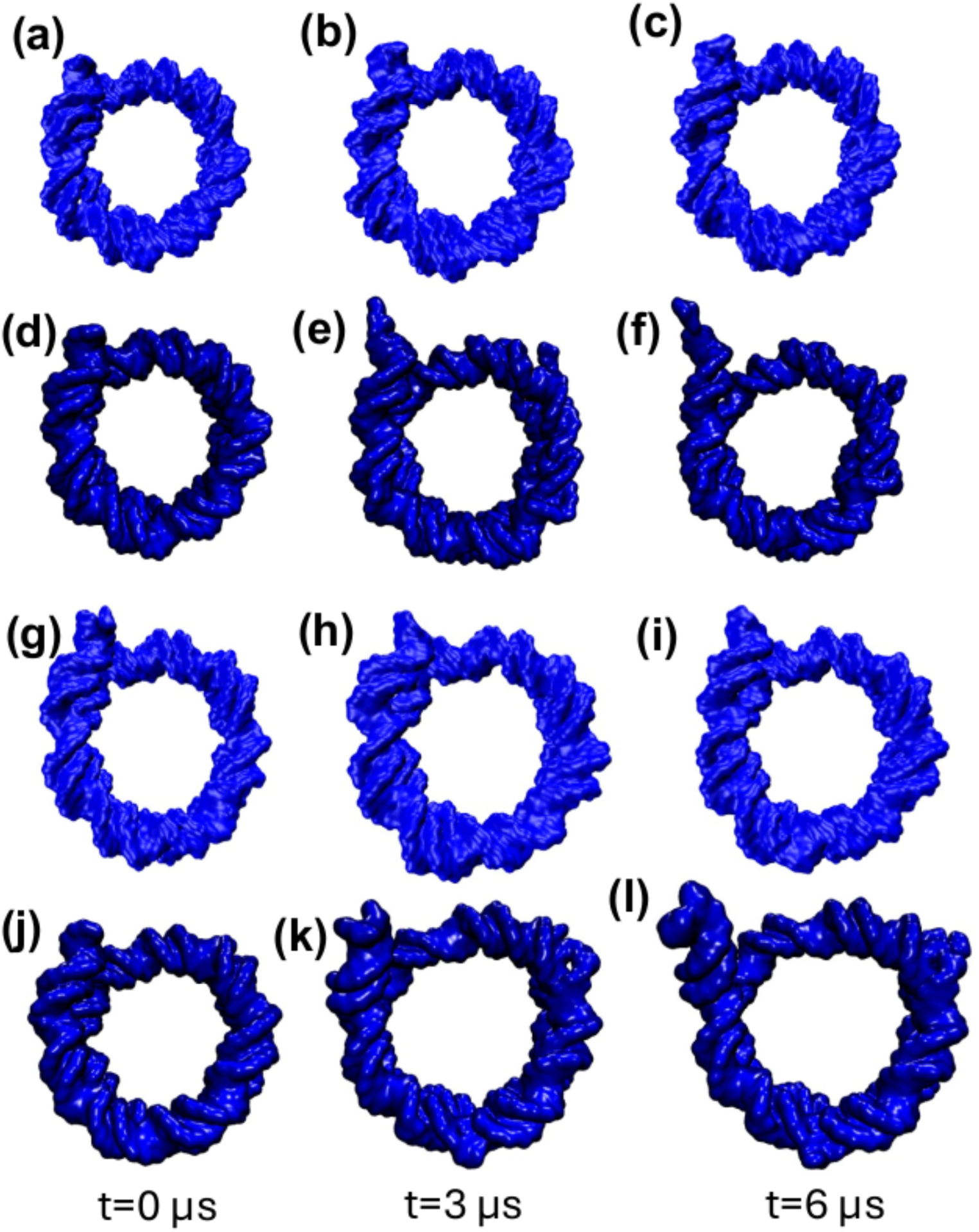
Snapshots illustrating motion of the nucleosomal DNA along the trajectory. The ASP DNA obtained from (a)-(c) atomistic simulation, (d)-(f) CG simulation. The Widom-601 sequence obtained from (g)-(i) atomistic and (j)-(l) CG simulations.

On the contrary, for the CG simulation (Fig. 6 d-f), the nucleosomal DNA shows substantial breathing motion at both t=3 µs (Fig. 6e) and t= 6 µs (Fig. 6f), respectively. We further check the breathing distance for both ends of the Widom-601 sequence. The histogram of breathing distance for End1 (601-L) (Fig. 5c) suggests a greater extent of breathing for the CG than the all-atom trajectories. The average value of breathing distance for the CG is 14.84 Å, while for AA, the average breathing distance is 3.24 Å. End2 (601-R) of the Widom-601 sequence shows similar behavior, i.e., a higher range of breathing distance for the CG than the AA (Fig. 5d). The average breathing distance for End2 is 6.84 Å. In contrast, for AA, the average value of breathing distance is 1.75 Å. Here, different ends also show differential breathing, like the ASP sequences. End1 (601-L) shows a higher distribution of breathing distances than End2, indicating asymmetric breathing. Fig. 6g-i shows the motion of DNA at different times for the Widom-601 sequence. The breathing motion is insignificant for the structures obtained from atomistic trajectory over the entire simulation of 6 µs. Meanwhile, structures obtained from CG simulations show substantial breathing motion (Fig. 6k-l). The SIRAH CG force field exhibits higher breathing than the AA simulations, with differential breathing motion for both ends of the DNA for both the ASP and Widom-601 sequences.

We further quantify the breathing calculating ΔR, which is displacement in average distance of each DNA base pair center represented in SHL notation over the simulated trajectory compared to the crystal structure in Fig. S6. We find a higher value of ΔR (nearly 10 Å) at SHL -7 for the ASP CG trajectories in one end, while the other has a lower value of ΔR (Fig. S6a). We did not find large values in ΔR in the atomistic simulation of ASP as in the CG trajectories, suggesting negligible breathing motion for both ends. The significant breathing motion is also visible for the Widom-601 sequence at both ends of DNA (Fig. S6b). However, the SHL +7 region (601-L) shows much higher breathing for the Widom-601 than the ASP sequence (ASP-L).

### 4.3 Principal Component Analysis (PCA) based on base-pair parameters

We further perform a conformational analysis of DNA based on the free energy landscape (FEL) obtained by projecting MD trajectories into the first two principal components, PC1 and PC2, for the DNA inter-base pair parameters (details in Methods). Fig. 7a shows the FEL for the CG trajectory of the ASP sequence, suggesting three different energy minima. We extract conformations from each minimum to better understand the conformation of the nucleosomal DNA. The end breathing distance for three different conformations from different clusters is substantially different. The extent of the distance for End1 is the maximum for the conformation from region ii (conformation 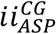), i.e., 22.79 Å. In contrast, from region iii (conformation 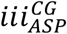), the value is lower, i.e., 1.04 Å (Fig. 7a). The DNA conformation from region i (conformation 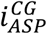) also shows a more significant breathing, i.e., 16.7 Å. The extent of breathing for End2 is lower than End1 for conformation from regions i and ii, but for region iii, the extent of breathing is higher. The conformation in region ii indicates inward movement as compared to crystal structure. The change in breathing distance for End2 is higher for conformations from Region ii, i.e., 11.93 Å, and from Region I, it is 5.31 Å. Overall, the FEL suggests conformations from different free energy minima show different levels of extent in breathing motion for both ends of the nucleosomal DNA. We find two different energy minima for the atomistic simulation for the ASP sequence (Fig. 7b). The conformation obtained from region i (conformation 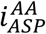) shows inward movement w.r.t crystal structure for both ends. End2 is showing a much larger extent than End1. The structure from Region ii (conformation 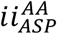) shows the opposite behavior, i.e., End1 shows a more significant extent of breathing motion than End2. For the atomistic simulations, the extent of breathing on both ends is lower than in the CG simulation, but the asymmetry in breathing distance between the two ends is maintained.

**Figure 7.**
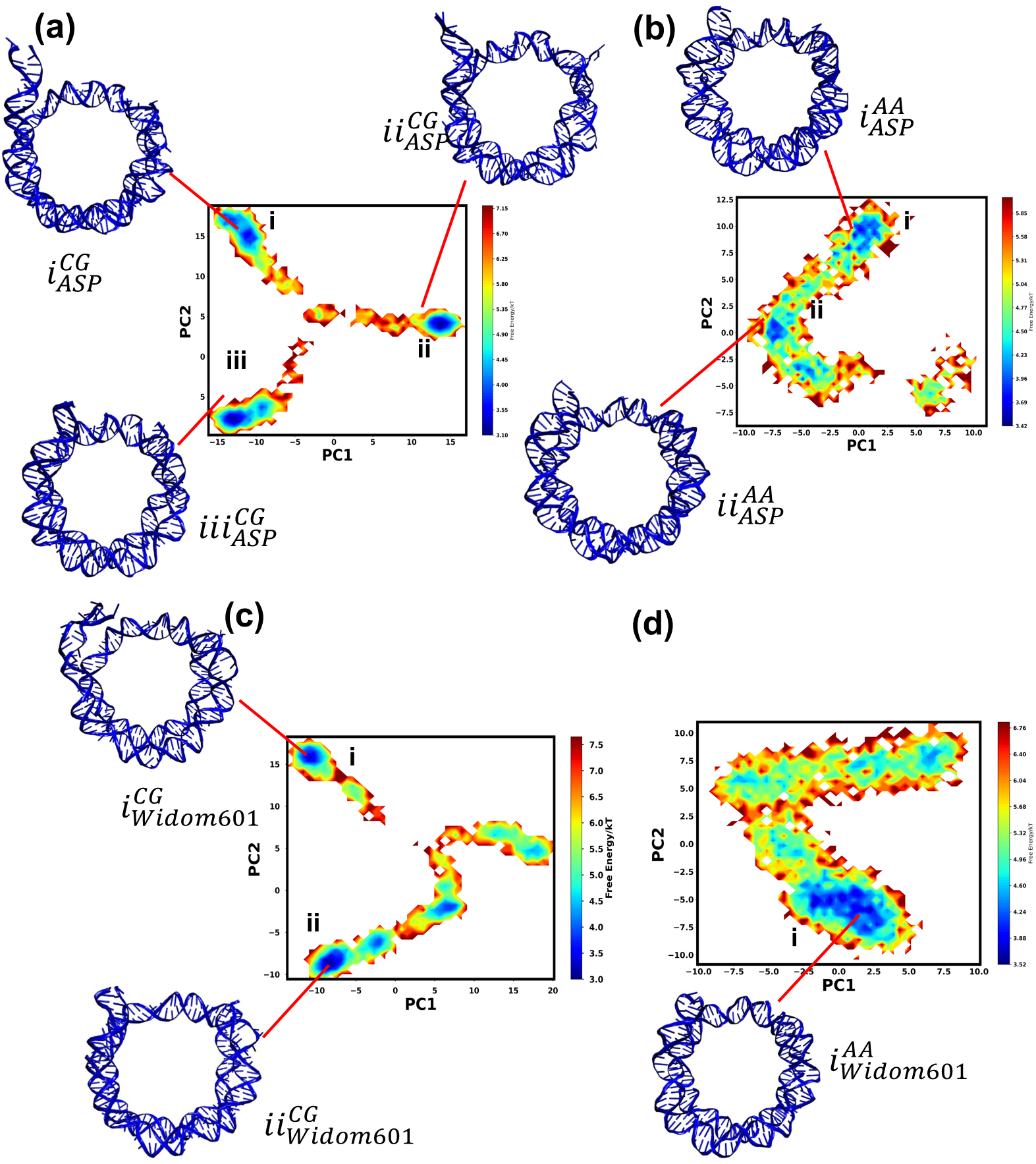
Principal component analysis (PCA) based on DNA inter base pair parameters. The Free energy landscape (FEL) based on PC1 and PC2 for (a) the coarse-grained trajectory, (b) atomistic simulation for the ASP sequence. The FEL for (c) the coarse-grained trajectory, (d) the atomistic simulation for the Widom-601 sequence. The energy minima are marked and structures with the minimum energy are shown.

Next, we extract conformations from the FEL for the Widom-601 sequence. Fig. 7c shows the FEL for the CG trajectories. We find two different minima (marked as i and ii) in PCA space. The DNA conformation from Region i (conformation 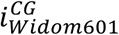) shows a breathing distance of 11.88 Å at End1 (601-L), while End2 (601-R) shows a breathing distance in the reverse direction of distance 1.72 Å. The conformation from Region ii (conformation 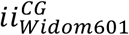) possesses a nearly equal breathing distance at End1 (601-L). It shows a distance of 11.49 Å, although End2 (601-R) shows a breathing distance of 2.51 Å. The atomistic simulation of Widom-601 indicates a single minimum (Fig. 7d, conformation 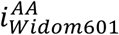). The breathing distance at both ends shows an inward breathing w.r.t the crystal structure. End1 (601-L) and End2 (601-R) show breathing distances of 3.87 Å and 0.81 Å, respectively. The atomistic simulation for Widom-601 shows a lower amount of breathing than the CG simulation within the simulated timescale. Still, the higher breathing distance of End1 (601-L) is maintained in both AA and CG simulations.

### 4.4 DNA repositioning around the histone core

To further understand nucleosomal dynamics, we probe nucleosomal DNA repositioning around the histone core using translocation (S_T_) and rotational (S_R_) order parameters (see analysis section). Fig. 8a shows a two-dimensional free energy plot for the ASP sequence as a function of S_T_ and S_R_, considering back-mapped CG trajectories using the SIRAH force field. The free energy surface suggests a strong tendency for rotational repositioning. However, translational repositioning is limited mostly within -0.4 to 0.4. The free energy minimum corresponds to S_T_ ≈ 0 with the minor groove towards the histone core. For the atomistic simulation (Fig. 8b), the free energy landscape indicates two distinct free energy minimums around S_T_ ≈ 0, with minor grooves towards the histone core. The free energy landscape for both S_T_ and S_R_ for the Widom-601 sequence shows multiple minima (Fig. 8c) for this sequence around positive values of S_T_. These free energy minima correspond to both S_R_ > 0 as well as S_R_ < 0. This suggests the minor groove is aligned towards and away from the histone core in the free energy minima. The FEL for the atomistic force field for the Widom-601 sequence (Fig. 8d) suggests two distinct minima in the free energy landscape. The difference with the CG counterpart is for AA, the energy minima correspond to S_T_ < 0, suggesting backward translocation of the nucleosomal DNA. Two distinct minima are observed at S_R_ > 0 and S_R_ < 0, suggesting a tendency to align minor grooves towards and away from the histone core. This behavior is similar to the ASP sequence (Fig. 8b). Overall, the result suggests that the CG force field can sample an extended range of possible states in the free energy landscape for both sequences, indicating multiple minima. In contrast, the AA force field restricts the system from exploring the available free energy landscape.

**Figure 8.**
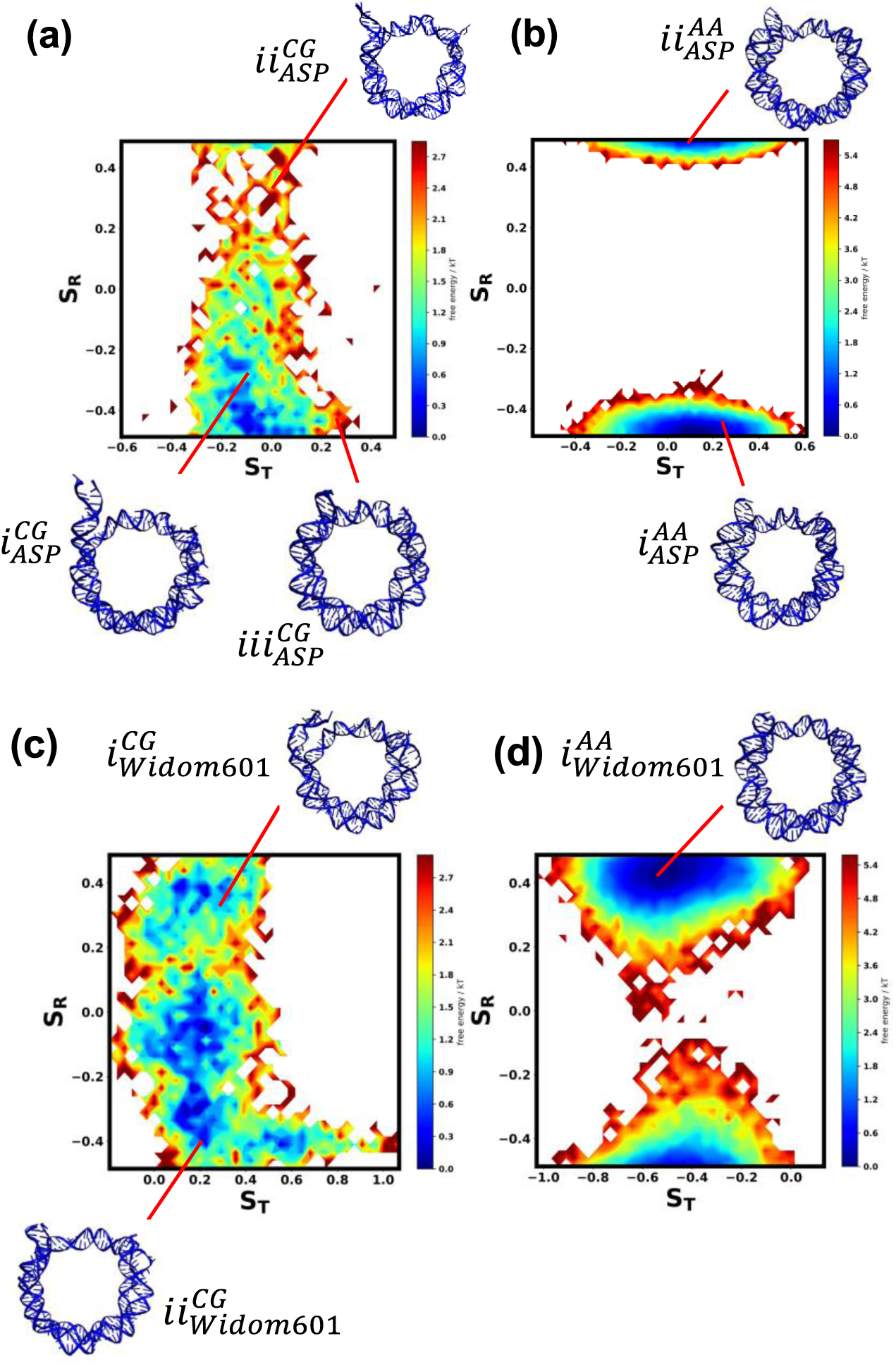
Free energy surface for DNA repositioning around histone based on (a) the coarse-grained trajectory, (b) the atomistic trajectory for the ASP sequence, (c) the coarse-grained trajectory, and (d) the atomistic trajectory of Widom-601 sequence.

## 5. Discussion

Overall, in this study, we focus on how the CG SIRAH force field can reproduce the conformations of nucleosomal DNA obtained using long-time molecular dynamics simulations using a state-of-the-art atomistic force field. Fig. 2b indicates a minimal difference in R_g_ for the ASP nucleosomal DNA between the CG and the AA model. The behavior of R_g_ is still preserved for the Widom-601 nucleosomal DNA (Fig. 2e). For the histone, we obtain a similar behavior, i.e., the difference in average R_g_ value between CG and AA trajectory is minimal. This indicates little deviation in R_g_ for both nucleosomal DNA and histone protein using the SIRAH ff compared to the AA forcefield. Next, we focus on various structural parameters, which mainly focus on the local geometry of the DNA. We characterize the groove width for the nucleosomal DNA. Fig. S3 indicates that average major and average minor width values do not deviate much between CG and AA trajectories. We compare inter-base pair parameters obtained from CG and AA trajectories for the ASP and Widom-601 DNA sequences. Most inter-base pair parameters show good agreement between CG and AA trajectories except for the roll inter-base pair parameter for both sequences. This study is consistent with earlier studies of DNA based on the SIRAH force field^76, 79^. The deviation for roll mainly occurs since the SIRAH ff is parametrized to reproduce the canonical B-form of DNA. In contrast, the OL15 ff is parametrized based on more extensive experimental structures^86^. We further compare various intra-base pair parameters of the nucleosomal DNA to understand better the structural similarity between AA and CG force fields. All intra-base pair parameters mainly show good similarity between AA and CG trajectories. However, propel and opening show a more significant deviation between CG and AA trajectories for both sequences. Despite some disparity in roll, propel, and opening order parameters between CG and AA trajectories, the SIRAH force field effectively captures most structural parameters. This motivates us to observe the extent of breathing motion for both End1 and End2 of the nucleosomal DNA. Neither sequence shows significant breathing motion within the simulated timescales using the atomistic force field in physiological salt concentration. However, the CG trajectory based on the SIRAH force field shows reasonable breathing motion for both End1 and End2 within the simulated time scale. This extent of breathing motion is observed for both sequences. For the ASP sequence, End1 (ASP-L) shows a more significant breathing motion than End2 (ASP-R). This result is consistent with earlier simulation results^23, 24^. Chakrabarty et al.^23^ showed that for the ASP sequence, a loop was formed at this same end (End1 (ASP-L)) as compared to End2 (ASP-R). This asymmetric breathing motion in our CG simulation also aligns with earlier experimental studies by Ngo and coworkers^12^. Using a single molecule optical trapping technique, they showed that one end interacts with the histone more strongly than the other as it is more flexible (601-L). Hence, a higher force is required to unwrap that end. Such an asymmetrical nature of DNA breathing is essential to understand as it might be a gene expression control factor affecting DNA exposure. In Khatua et al.,^24^ we find a similar result for the ASP sequence based on our 12 µs simulation in high salt conditions. However, we find much larger breathing in the case of the Widom-601 sequence. End 2 (601-R) shows more extensive breathing than End 1 (601-L); furthermore, the overall breathing motion is higher in the Widom-601 sequence than in the ASP sequence. Conversely, in our CG study for both sequences, we found no sequence-specific bias regarding breathing distance; indeed, End1(601-L) shows more significant breathing motion than End2 (601- R).

We next conduct further analysis of breathing distance in DNA conformations obtained after performing PCA on the DNA base pair parameters. We identify both outward and inward breathing motion w.r.t crystal structure for both sequences at both Ends. Multiple minima with higher breathing distances have been observed for CG trajectories compared to atomistic simulation.

Hence, the SIRAH CG force field can efficiently sample multiple minima relative to the atomistic force field. Next, we investigate the repositioning of the nucleosomal DNA around histone. To understand the DNA repositioning around a histone, we calculate two additional order parameters, i.e., S_T_ and S_R_. We show the free energy surface based on both S_T_ and S_R_. The SIRAH CG force field can sample multiple minima of the free energy landscape, while the AA force field shows restricted dynamics within specific regions of the free energy landscape. This result is consistent with earlier results of the FEL based on PCA of inter-base pair parameters.

We further elucidate the DNA repositioning mechanism from these free energy surfaces. In the free energy surface based on the base pair parameter, we find different free energy minima at different positions of the free energy surface (Fig. 7a and 7c). We next identify the conformations at different minima and identified those conformations on the free energy surface obtained using translation (S_T_) and rotation (S_R_) order parameters (Fig. 8). Different conformations are marked on the free energy surface on Fig. 8. For the ASP sequence, one conformation belongs to energy minima for the CG trajectory (conformation 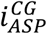) (Fig. 8a). For the ASP atomistic simulation, we find two conformations at two different energy minima (Fig. 8b) at two different regions, suggesting restricted sampling in those energy minima. For the Widom-601 atomistic simulation, we find similar behavior, i.e., two distinct conformations at two different energy minima (Fig. 8d). For the case of the Widom-601 sequence, we find two distinct conformations at two different energy minima (Fig. 8c) along with possible multiple paths between those conformations. Conformation 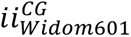 belongs to a region where S_R_<0 while the other conformation 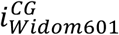 belongs to the S_R_ > 0 region. We identify a minimum free energy path between those two conformations (Fig. 9) using the “String Method.” We plot the minimum free energy path between the two conformations in Fig. 9. Fig. 9 also shows the structures over the free energy path, suggesting that the breathing motion of DNA is accompanied by DNA rotation around the histone core. In contrast, Lequieu et al.^60^ report the DNA repositioning mechanism for Widom-601 is almost independent of rotational position. This difference in mechanism suggests a more careful analysis of the local twisting of DNA in particular SHL regions may be necessary to elucidate the mechanism further. For example, Armeev et al. ^26^ have observed twist defects, etc. We have also observed twisting in particular regions^24^ of the DNA at high salt concentrations. Notably, we have not observed loop propagation for this set of simulations as we have observed at high salt concentrations. Both loop propagation^106–109^ and twist diffusion^110–112^ have been reported. Some experimental evidence supports both mechanisms^110–114^.

**Figure 9.**
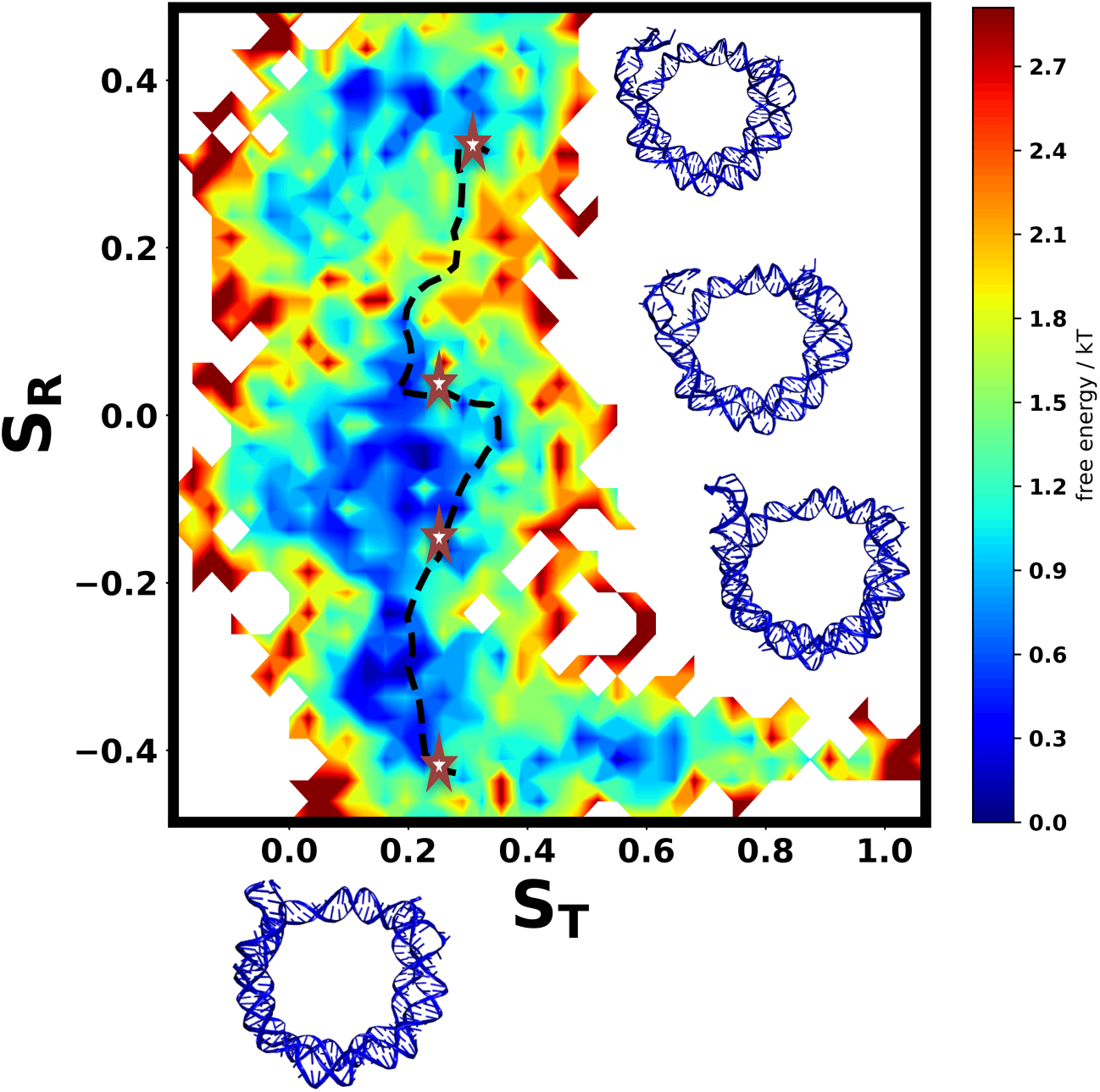
The minimum free energy path corresponding to DNA rotation for the Widom-601 sequence. The star corresponds to different conformational states of the DNA obtained from energy minima of free energy landscape based on PCA of the inter base pair parameters

## 6. Conclusions

In summary, the simulations presented here explore nucleosome dynamics at physiological salt concentration for both atomistic and SIRAH CG force fields at the time scale of microseconds. Simulation of the two nucleosome systems containing different DNA sequences, the ASP and the Widom-601 sequence, using the SIRAH CG force field, capture major conformations of nucleosomal DNA. We obtain a greater extent of breathing motion of both Ends of the DNA in CG simulations relative to atomistic simulations. Principal component analysis based on DNA dinucleotide base pair parameters aids in identifying multiple minima in the free energy landscape for both the CG and atomistic force fields. The SIRAH CG force field explores multiple minima relative to atomistic trajectories. CG simulations preserve the asymmetric motion observed for the DNA ends. Next, we construct a minimum energy path based on the free energy landscape for nucleosome repositioning. For this set of simulations, we find that the Widom-601 sequence involves rotational repositioning. We hypothesize that the transition between different states can be probed using Markov state models (MSMs)^115, 116^. This approach can provide information based on kinetic exchange between different conformational states of nucleosomal DNA. The SIRAH CG forcefield has significant potential to address the dynamics of larger protein-DNA complexes like tetranucleosomes^28^ or address the shift in DNA repositioning with the binding of transcription factors ^117^ and chromatin remodelers^118^. We note that the histone tail parameters and their interaction with the DNA may need to be subtly altered to better match conformational fluctuations of the tails in atomistic simulations and order parameters of the tails that can be observed via NMR spectroscopy.

## Supplementary Material

See Supplementary Material Fig. S1-S6. Fig. S1 includes schematics of Inter-base pair parameters, Intra-base pair parameters, translocation, and rotational movement of nucleosomal DNA around histone. Fig. S2 includes time evolution and histogram of R_g_ of the histone and secondary structure percentage over CG and AA trajectories. Fig.S3 contains distributions of the DNA’s major and minor groove widths over both AA and CG trajectories. The histogram of intra0base pair parameters for both sequences is in Fig.S4-S5 for both atomistic and CG trajectories. Fig.S6 includes the change in the average distance of each DNA base pair center represented in SHL notation over the simulated trajectory compared to the crystal structure for both sequences. Supplementary Movie 1 and 2 contain the SIRAH CG trajectories for both sequences.

## Acknowledgements

This work was supported by NIH through Grant 1R15GM146228-01. Anton 2 computer time was provided by the Pittsburgh Supercomputing Center (PSC) through Grant R01GM116961 from the National Institutes of Health. The Anton 2 machine at PSC was generously made available by D.E. Shaw Research. We thank Prof. Sergio Pantano for their comments and help with SIRAH.

## Data Availability Statement

Analysis codes are available on https://github.com/CUNY-CSI-Loverde-Laboratory/GhoshMoulick_2024-. Trajectories are available on https://zenodo.org/records/14033991.

## References

(1) Luger, K.; Dechassa, M. L.; Tremethick, D. J. New insights into nucleosome and chromatin structure: An ordered state or a disordered affair? Nature reviews Molecular cell biology 2012, 13 (7), 436–447.

(2) McGinty, R. K.; Tan, S. Nucleosome structure and function. Chemical reviews 2015, 115 (6), 2255–2273.

(3) Kim, K.-D. Potential roles of condensin in genome organization and beyond in fission yeast. Journal of Microbiology 2021, 59 (5), 449–459.

(4) Kornberg, R. D.; Lorch, Y. Twenty-five years of the nucleosome, fundamental particle of the eukaryote chromosome. Cell 1999, 98 (3), 285–294.

(5) Segal, E.; Fondufe-Mittendorf, Y.; Chen, L.; Thåström, A.; Field, Y.; Moore, I. K.; Wang, J.-P. Z.; Widom, J. A genomic code for nucleosome positioning. Nature 2006, 442 (7104), 772–778.

(6) Müller, M. M.; Muir, T. W. Histones: At the crossroads of peptide and protein chemistry. Chemical reviews 2015, 115 (6), 2296–2349.

(7) Parmar, J. J.; Padinhateeri, R. Nucleosome positioning and chromatin organization. Current Opinion in Structural Biology 2020, 64, 111–118.

(8) Shaytan, A. K.; Armeev, G. A.; Goncearenco, A.; Zhurkin, V. B.; Landsman, D.; Panchenko, A. R. Coupling between histone conformations and DNA geometry in nucleosomes on a microsecond timescale: Atomistic insights into nucleosome functions. Journal of molecular biology 2016, 428 (1), 221–237.

(9) Li, Z.; Kono, H. Distinct roles of histone h3 and h2a tails in nucleosome stability. Scientific reports 2016, 6 (1), 31437.

(10) Gansen, A.; Hauger, F.; Toth, K.; Langowski, J. Single-pair fluorescence resonance energy transfer of nucleosomes in free diffusion: Optimizing stability and resolution of subpopulations. Analytical biochemistry 2007, 368 (2), 193–204.

(11) Chen, Y.; Tokuda, J. M.; Topping, T.; Meisburger, S. P.; Pabit, S. A.; Gloss, L. M.; Pollack, L. Asymmetric unwrapping of nucleosomal DNA propagates asymmetric opening and dissociation of the histone core. Proceedings of the National Academy of Sciences 2017, 114 (2), 334–339.

(12) Ngo, T. T.; Zhang, Q.; Zhou, R.; Yodh, J. G.; Ha, T. Asymmetric unwrapping of nucleosomes under tension directed by DNA local flexibility. Cell 2015, 160 (6), 1135–1144.

(13) Meersseman, G.; Pennings, S.; Bradbury, E. M. Mobile nucleosomes--a general behavior. The EMBO journal 1992, 11 (8), 2951–2959.

(14) Pennings, S.; Meersseman, G.; Bradbury, E. M. Mobility of positioned nucleosomes on 5 s rdna. Journal of molecular biology 1991, 220 (1), 101–110.

(15) Flaus, A.; Richmond, T. J. Positioning and stability of nucleosomes on mmtv 3′ ltr sequences. Journal of molecular biology 1998, 275 (3), 427–441.

(16) Materese, C. K.; Savelyev, A.; Papoian, G. A. Counterion atmosphere and hydration patterns near a nucleosome core particle. Journal of the American Chemical Society 2009, 131 (41), 15005–15013.

(17) Erler, J.; Zhang, R.; Petridis, L.; Cheng, X.; Smith, J. C.; Langowski, J. The role of histone tails in the nucleosome: A computational study. Biophysical journal 2014, 107 (12), 2911–2922.

(18) Morrison, E. A.; Bowerman, S.; Sylvers, K. L.; Wereszczynski, J.; Musselman, C. A. The conformation of the histone h3 tail inhibits association of the bptf phd finger with the nucleosome. Elife 2018, 7, e31481.

(19) Huertas, J.; Cojocaru, V. Breaths, twists, and turns of atomistic nucleosomes. Journal of molecular biology 2021, 433 (6), 166744.

(20) Ettig, R.; Kepper, N.; Stehr, R.; Wedemann, G.; Rippe, K. Dissecting DNA-histone interactions in the nucleosome by molecular dynamics simulations of DNA unwrapping. Biophysical journal 2011, 101 (8), 1999–2008.

(21) Rychkov, G. N.; Ilatovskiy, A. V.; Nazarov, I. B.; Shvetsov, A. V.; Lebedev, D. V.; Konev, A. Y.; Isaev-Ivanov, V. V.; Onufriev, A. V. Partially assembled nucleosome structures at atomic detail. Biophysical journal 2017, 112 (3), 460–472.

(22) Zhang, B.; Zheng, W.; Papoian, G. A.; Wolynes, P. G. Exploring the free energy landscape of nucleosomes. Journal of the American Chemical Society 2016, 138 (26), 8126–8133.

(23) Chakraborty, K.; Loverde, S. M. Asymmetric breathing motions of nucleosomal DNA and the role of histone tails. The Journal of Chemical Physics 2017, 147 (6).

(24) Khatua, P.; Tang, P. K.; Ghosh Moulick, A.; Patel, R.; Manandhar, A.; Loverde, S. M. Sequence dependence in nucleosome dynamics. The Journal of Physical Chemistry B 2024, 128 (13), 3090–3101.

(25) Chakraborty, K.; Kang, M.; Loverde, S. M. Molecular mechanism for the role of the h2a and h2b histone tails in nucleosome repositioning. The Journal of Physical Chemistry B 2018, 122 (50), 11827–11840.

(26) Armeev, G. A.; Kniazeva, A. S.; Komarova, G. A.; Kirpichnikov, M. P.; Shaytan, A. K. Histone dynamics mediate DNA unwrapping and sliding in nucleosomes. Nature communications 2021, 12 (1), 2387.

(27) Winogradoff, D.; Aksimentiev, A. Molecular mechanism of spontaneous nucleosome unraveling. Journal of molecular biology 2019, 431 (2), 323–335.

(28) Ding, X. Q.; Lin, X. C.; Zhang, B. Stability and folding pathways of tetra-nucleosome from six-dimensional free energy surface. Nature Communications 2021, 12 (1).

(29) Farr, S. E.; Woods, E. J.; Joseph, J. A.; Garaizar, A.; Collepardo-Guevara, R. Nucleosome plasticity is a critical element of chromatin liquid–liquid phase separation and multivalent nucleosome interactions. Nature communications 2021, 12 (1), 2883.

(30) Yoo, J.; Winogradoff, D.; Aksimentiev, A. Molecular dynamics simulations of DNA–DNA and DNA–protein interactions. Current Opinion in Structural Biology 2020, 64, 88–96.

(31) Pérez, A.; Marchán, I.; Svozil, D.; Sponer, J.; Cheatham, T. E.; Laughton, C. A.; Orozco, M. Refinement of the amber force field for nucleic acids: Improving the description of α/γ conformers. Biophysical journal 2007, 92 (11), 3817–3829.

(32) Ivani, I.; Dans, P. D.; Noy, A.; Pérez, A.; Faustino, I.; Hospital, A.; Walther, J.; Andrio, P.; Goñi, R.; Balaceanu, A. Parmbsc1: A refined force field for DNA simulations. Nature methods 2016, 13 (1), 55–58.

(33) Zgarbová, M.; Sponer, J.; Otyepka, M.; Cheatham III, T. E.; Galindo-Murillo, R.; Jurecka, P. Refinement of the sugar–phosphate backbone torsion beta for amber force fields improves the description of z-and b-DNA. Journal of chemical theory and computation 2015, 11 (12), 5723–5736.

(34) Denning, E. J.; Priyakumar, U. D.; Nilsson, L.; Mackerell Jr, A. D. Impact of 2′-hydroxyl sampling on the conformational properties of rna: Update of the charmm all-atom additive force field for rna. Journal of computational chemistry 2011, 32 (9), 1929–1943.

(35) Hart, K.; Foloppe, N.; Baker, C. M.; Denning, E. J.; Nilsson, L.; MacKerell Jr, A. D. Optimization of the charmm additive force field for DNA: Improved treatment of the bi/bii conformational equilibrium. Journal of chemical theory and computation 2012, 8 (1), 348–362.

(36) Cornell, W. D.; Cieplak, P.; Bayly, C. I.; Gould, I. R.; Merz, K. M.; Ferguson, D. M.; Spellmeyer, D. C.; Fox, T.; Caldwell, J. W.; Kollman, P. A. A second generation force field for the simulation of proteins, nucleic acids, and organic molecules. Journal of the American Chemical Society 1995, 117 (19), 5179–5197.

(37) Zgarbová, M.; Otyepka, M.; Sponer, J.; Mladek, A.; Banas, P.; Cheatham III, T. E.; Jurecka, P. Refinement of the cornell et al. Nucleic acids force field based on reference quantum chemical calculations of glycosidic torsion profiles. Journal of chemical theory and computation 2011, 7 (9), 2886–2902.

(38) Galindo-Murillo, R.; Robertson, J. C.; Zgarbova, M.; Sponer, J.; Otyepka, M.; Jurecka, P.; Cheatham III, T. E. Assessing the current state of amber force field modifications for DNA. Journal of chemical theory and computation 2016, 12 (8), 4114–4127.

(39) Love, O.; Galindo-Murillo, R.; Zgarbová, M.; Šponer, J. i.; Jurečka, P.; Cheatham III, T. E. Assessing the current state of amber force field modifications for DNA─ 2023 edition. Journal of Chemical Theory and Computation 2023, 19 (13), 4299–4307.

(40) Minhas, V.; Sun, T.; Mirzoev, A.; Korolev, N.; Lyubartsev, A. P.; Nordenskiöld, L. Modeling DNA flexibility: Comparison of force fields from atomistic to multiscale levels. The Journal of Physical Chemistry B 2019, 124 (1), 38–49.

(41) Tucker, M. R.; Piana, S.; Tan, D.; LeVine, M. V.; Shaw, D. E. Development of force field parameters for the simulation of single-and double-stranded DNA molecules and DNA–protein complexes. The Journal of Physical Chemistry B 2022, 126 (24), 4442–4457.

(42) Wei, S. J.; Falk, S. J.; Black, B. E.; Lee, T. H. A novel hybrid single molecule approach reveals spontaneous DNA motion in the nucleosome. Nucleic Acids Research 2015, 43 (17), E111–U148.

(43) Bilokapic, S.; Strauss, M.; Halic, M. Structural rearrangements of the histone octamer translocate DNA. Nature Communications 2018, 9 (1), 1330.

(44) Bowman, G. D.; Poirier, M. G. Post-translational modifications of histones that influence nucleosome dynamics. Chemical Reviews 2015, 115 (6), 2274–2295.

(45) Patel, R.; Onyema, A.; Tang, P. K.; Loverde, S. M. Conformational dynamics of the nucleosomal histone h2b tails revealed by molecular dynamics simulations. Journal of Chemical Information and Modeling 2024.

(46) Ozer, G.; Luque, A.; Schlick, T. The chromatin fiber: Multiscale problems and approaches. Current Opinion in Structural Biology 2015, 31, 124–139.

(47) Hyeon, C.; Thirumalai, D. Capturing the essence of folding and functions of biomolecules using coarse-grained models. Nature communications 2011, 2 (1), 487.

(48) Reddy, G.; Thirumalai, D. Asymmetry in histone rotation in forced unwrapping and force quench rewrapping in a nucleosome. Nucleic acids research 2021, 49 (9), 4907–4918.

(49) Lequieu, J.; Córdoba, A.; Schwartz, D. C.; de Pablo, J. J. Tension-dependent free energies of nucleosome unwrapping. ACS central science 2016, 2 (9), 660–666.

(50) Sun, T.; Minhas, V.; Mirzoev, A.; Korolev, N.; Lyubartsev, A. P.; Nordenskiöld, L. A bottom-up coarse-grained model for nucleosome–nucleosome interactions with explicit ions. Journal of Chemical Theory and Computation 2022, 18 (6), 3948–3960.

(51) Chakraborty, D.; Mondal, B.; Thirumalai, D. Brewing coffee: A sequence-specific coarse-grained energy function for simulations of DNA−protein complexes. Journal of Chemical Theory and Computation 2024, 20 (3), 1398–1413.

(52) Li, Z.; Portillo-Ledesma, S.; Schlick, T. Brownian dynamics simulations of mesoscale chromatin fibers. Biophysical journal 2023, 122 (14), 2884–2897.

(53) Beard, D. A.; Schlick, T. Computational modeling predicts the structure and dynamics of chromatin fiber. Structure 2001, 9 (2), 105–114.

(54) Zhang, Q.; Beard, D. A.; Schlick, T. Constructing irregular surfaces to enclose macromolecular complexes for mesoscale modeling using the discrete surface charge optimization (disco) algorithm. Journal of computational chemistry 2003, 24 (16), 2063–2074.

(55) Collepardo-Guevara, R.; Schlick, T. Chromatin fiber polymorphism triggered by variations of DNA linker lengths. Proceedings of the National Academy of Sciences 2014, 111 (22), 8061–8066.

(56) Arya, G.; Schlick, T. Role of histone tails in chromatin folding revealed by a mesoscopic oligonucleosome model. Proceedings of the National Academy of Sciences 2006, 103 (44), 16236–16241.

(57) Perišić, O.; Portillo-Ledesma, S.; Schlick, T. Sensitive effect of linker histone binding mode and subtype on chromatin condensation. Nucleic Acids Research 2019, 47 (10), 4948–4957.

(58) Davtyan, A.; Schafer, N. P.; Zheng, W.; Clementi, C.; Wolynes, P. G.; Papoian, G. A. Awsem-md: Protein structure prediction using coarse-grained physical potentials and bioinformatically based local structure biasing. The Journal of Physical Chemistry B 2012, 116 (29), 8494–8503.

(59) Hinckley, D. M.; Freeman, G. S.; Whitmer, J. K.; De Pablo, J. J. An experimentally-informed coarse-grained 3-site-per-nucleotide model of DNA: Structure, thermodynamics, and dynamics of hybridization. The Journal of chemical physics 2013, 139 (14).

(60) Lequieu, J.; Schwartz, D. C.; de Pablo, J. J. In silico evidence for sequence-dependent nucleosome sliding. Proceedings of the National Academy of Sciences 2017, 114 (44), E9197–E9205.

(61) Niina, T.; Brandani, G. B.; Tan, C.; Takada, S. Sequence-dependent nucleosome sliding in rotation-coupled and uncoupled modes revealed by molecular simulations. PLoS computational biology 2017, 13 (12), e1005880.

(62) Brandani, G. B.; Niina, T.; Tan, C.; Takada, S. DNA sliding in nucleosomes via twist defect propagation revealed by molecular simulations. Nucleic acids research 2018, 46 (6), 2788–2801.

(63) Nagae, F.; Brandani, G. B.; Takada, S.; Terakawa, T. The lane-switch mechanism for nucleosome repositioning by DNA translocase. Nucleic Acids Research 2021, 49 (16), 9066–9076.

(64) Brandner, A.; Schüller, A.; Melo, F.; Pantano, S. Exploring DNA dynamics within oligonucleosomes with coarse-grained simulations: Sirah force field extension for protein-DNA complexes. Biochemical and biophysical research communications 2018, 498 (2), 319–326.

(65) Honorato, R. V.; Roel-Touris, J.; Bonvin, A. M. Martini-based protein-DNA coarse-grained haddocking. Frontiers in molecular biosciences 2019, 6, 102.

(66) Borges-Araújo, L.; Patmanidis, I.; Singh, A. P.; Santos, L. H.; Sieradzan, A. K.; Vanni, S.; Czaplewski, C.; Pantano, S.; Shinoda, W.; Monticelli, L. Pragmatic coarse-graining of proteins: Models and applications. Journal of Chemical Theory and Computation 2023, 19 (20), 7112–7135.

(67) Borges-Araujo, L.; Patmanidis, I.; Singh, A. P.; Santos, L. H. S.; Sieradzan, A. K.; Vanni, S.; Czaplewski, C.; Pantano, S.; Shinoda, W.; Monticelli, L.;, et al. Pragmatic coarse-graining of proteins: Models and applications. Journal of Chemical Theory and Computation 2023, 19 (20), 7112–7135.

(68) Uusitalo, J. J.; Ingólfsson, H. I.; Akhshi, P.; Tieleman, D. P.; Marrink, S. J. Martini coarse-grained force field: Extension to DNA. Journal of chemical theory and computation 2015, 11 (8), 3932–3945.

(69) Klein, F.; Soñora, M.; Santos, L. H.; Frigini, E. N.; Ballesteros-Casallas, A.; Machado, M. R.; Pantano, S. The sirah force field: A suite for simulations of complex biological systems at the coarse-grained and multiscale levels. Journal of structural biology 2023, 215 (3), 107985.

(70) Klein, F.; Soñora, M.; Santos, L. H.; Frigini, E. N.; Ballesteros-Casallas, A.; Machado, M. R.; Pantano, S. The sirah force field: A suite for simulations of complex biological systems at the coarse-grained and multiscale levels. Journal of structural biology 2023, 107985.

(71) Darré, L.; Machado, M. R.; Brandner, A. F.; González, H. C.; Ferreira, S.; Pantano, S. Sirah: A structurally unbiased coarse-grained force field for proteins with aqueous solvation and long-range electrostatics. Journal of chemical theory and computation 2015, 11 (2), 723–739.

(72) Machado, M. R.; Barrera, E. E.; Klein, F.; Sóñora, M.; Silva, S.; Pantano, S. The sirah 2.0 force field: Altius, fortius, citius. Journal of chemical theory and computation 2019, 15 (4), 2719–2733.

(73) Garay, P. G.; Barrera, E. E.; Pantano, S. Post-translational modifications at the coarse-grained level with the sirah force field. Journal of Chemical Information and Modeling 2019, 60 (2), 964–973.

(74) Klein, F.; Cáceres, D.; Carrasco, M. A.; Tapia, J. C.; Caballero, J.; Alzate-Morales, J.; Pantano, S. Coarse-grained parameters for divalent cations within the sirah force field. Journal of Chemical Information and Modeling 2020, 60 (8), 3935–3943.

(75) Barrera, E. E.; Machado, M. R.; Pantano, S. Fat sirah: Coarse-grained phospholipids to explore membrane–protein dynamics. Journal of Chemical Theory and Computation 2019, 15 (10), 5674–5688.

(76) Dans, P. D.; Zeida, A.; Machado, M. R.; Pantano, S. A coarse grained model for atomic-detailed DNA simulations with explicit electrostatics. Journal of Chemical Theory and Computation 2010, 6 (5), 1711–1725.

(77) Klein, F.; Barrera, E. E.; Pantano, S. Assessing sirah’s capability to simulate intrinsically disordered proteins and peptides. Journal of Chemical Theory and Computation 2021, 17 (2), 599–604.

(78) Machado, M. R.; Pantano, S. Exploring laci–DNA dynamics by multiscale simulations using the sirah force field. Journal of Chemical Theory and Computation 2015, 11 (10), 5012–5023.

(79) Dans, P. D.; Darré, L.; Machado, M. R.; Zeida, A.; Brandner, A. F.; Pantano, S. Assessing the accuracy of the sirah force field to model DNA at coarse grain level. In Advances in Bioinformatics and Computational Biology: 8th Brazilian Symposium on Bioinformatics, BSB 2013, Recife, Brazil, November 3-7, 2013, Proceedings 8, 2013; Springer: pp 71–81.

(80) Lequieu, J.; Schwartz, D. C.; de Pablo, J. J. In silico evidence for sequence-dependent nucleosome sliding. Proc Natl Acad Sci U S A 2017, 114 (44), E9197–e9205.

(81) Davey, C. A.; Sargent, D. F.; Luger, K.; Maeder, A. W.; Richmond, T. J. Solvent mediated interactions in the structure of the nucleosome core particle at 1.9å resolution††we dedicate this paper to the memory of max perutz who was particularly inspirational and supportive to t.J.R. In the early stages of this study. Journal of Molecular Biology 2002, 319 (5), 1097–1113.

(82) Vasudevan, D.; Chua, E. Y. D.; Davey, C. A. Crystal structures of nucleosome core particles containing the ‘601’ strong positioning sequence. Journal of Molecular Biology 2010, 403 (1), 1–10.

(83) Jacobson, M. P.; Friesner, R. A.; Xiang, Z.; Honig, B. On the role of the crystal environment in determining protein side-chain conformations. J Mol Biol 2002, 320 (3), 597–608.

(84) Jacobson, M. P.; Pincus, D. L.; Rapp, C. S.; Day, T. J.; Honig, B.; Shaw, D. E.; Friesner, R. A. A hierarchical approach to all-atom protein loop prediction. Proteins 2004, 55 (2), 351–367.

(85) Tian, C.; Kasavajhala, K.; Belfon, K. A. A.; Raguette, L.; Huang, H.; Migues, A. N.; Bickel, J.; Wang, Y.; Pincay, J.; Wu, Q.;, et al. Ff19sb: Amino-acid-specific protein backbone parameters trained against quantum mechanics energy surfaces in solution. J Chem Theory Comput 2020, 16 (1), 528–552.

(86) Zgarbová, M.; Šponer, J.; Otyepka, M.; Cheatham, T. E., 3rd; Galindo-Murillo, R.; Jurečka, P. Refinement of the sugar-phosphate backbone torsion beta for amber force fields improves the description of z- and b-DNA. J Chem Theory Comput 2015, 11 (12), 5723–5736.

(87) Izadi, S.; Anandakrishnan, R.; Onufriev, A. V. Building water models: A different approach. J Phys Chem Lett 2014, 5 (21), 3863–3871.

(88) Joung, I. S.; Cheatham, T. E., 3rd. Determination of alkali and halide monovalent ion parameters for use in explicitly solvated biomolecular simulations. J Phys Chem B 2008, 112 (30), 9020–9041.

(89) Li, Z.; Song, L. F.; Li, P.; Merz, K. M., Jr. Systematic parametrization of divalent metal ions for the opc3, opc, tip3p-fb, and tip4p-fb water models. J Chem Theory Comput 2020, 16 (7), 4429–4442.

(90) Kulkarni, M.; Yang, C.; Pak, Y. Refined alkali metal ion parameters for the opc water model. Bulletin of the Korean Chemical Society 2018, 39 (8), 931–935.

(91) Case, D. A.; Cheatham III, T. E.; Darden, T.; Gohlke, H.; Luo, R.; Merz Jr, K. M.; Onufriev, A.; Simmerling, C.; Wang, B.; Woods, R. J. The amber biomolecular simulation programs. Journal of computational chemistry 2005, 26 (16), 1668–1688.

(92) Andersen, H. C. Rattle: A “velocity” version of the shake algorithm for molecular dynamics calculations. Journal of computational Physics 1983, 52 (1), 24–34.

(93) Shaw, D. E.; Grossman, J.; Bank, J. A.; Batson, B.; Butts, J. A.; Chao, J. C.; Deneroff, M. M.; Dror, R. O.; Even, A.; Fenton, C. H. Anton 2: Raising the bar for performance and programmability in a special-purpose molecular dynamics supercomputer. In SC’14: Proceedings of the International Conference for High Performance Computing, Networking, Storage and Analysis, 2014; IEEE: pp 41–53.

(94) Van Der Spoel, D.; Lindahl, E.; Hess, B.; Groenhof, G.; Mark, A. E.; Berendsen, H. J. Gromacs: Fast, flexible, and free. Journal of computational chemistry 2005, 26 (16), 1701–1718.

(95) Dolinsky, T. J.; Nielsen, J. E.; McCammon, J. A.; Baker, N. A. Pdb2pqr: An automated pipeline for the setup of poisson–boltzmann electrostatics calculations. Nucleic acids research 2004, 32 (suppl_2), W665-W667.

(96) Darré, L.; Machado, M. R.; Dans, P. D.; Herrera, F. E.; Pantano, S. Another coarse grain model for aqueous solvation: Wat four? Journal of Chemical Theory and Computation 2010, 6 (12), 3793–3807.

(97) Bussi, G.; Donadio, D.; Parrinello, M. Canonical sampling through velocity rescaling. The Journal of chemical physics 2007, 126 (1).

(98) Machado, M. R.; Pantano, S. Sirah tools: Mapping, backmapping and visualization of coarse-grained models. Bioinformatics 2016, 32 (10), 1568–1570.

(99) Parsons, J.; Holmes, J. B.; Rojas, J. M.; Tsai, J.; Strauss, C. E. M. Practical conversion from torsion space to cartesian space for in silico protein synthesis. Journal of Computational Chemistry 2005, 26 (10), 1063–1068.

(100) Maier, J. A.; Martinez, C.; Kasavajhala, K.; Wickstrom, L.; Hauser, K. E.; Simmerling, C. Ff14sb: Improving the accuracy of protein side chain and backbone parameters from ff99sb. Journal of chemical theory and computation 2015, 11 (8), 3696–3713.

(101) Case, D. A.; Aktulga, H. M.; Belfon, K.; Cerutti, D. S.; Cisneros, G. A.; Cruzeiro, V. W. D.; Forouzesh, N.; Giese, T. J.; Götz, A. W.; Gohlke, H. Ambertools. Journal of chemical information and modeling 2023, 63 (20), 6183–6191.

(102) Frishman, D.; Argos, P. Knowledge - based protein secondary structure assignment. *Proteins: Structure*, Function, and Bioinformatics 1995, 23 (4), 566–579.

(103) Lavery, R.; Moakher, M.; Maddocks, J. H.; Petkeviciute, D.; Zakrzewska, K. Conformational analysis of nucleic acids revisited: Curves+. Nucleic acids research 2009, 37 (17), 5917–5929.

(104) E, W.; Ren, W.; Vanden-Eijnden, E. String method for the study of rare events. Physical Review B 2002, 66 (5), 052301.

(105) Qiu, C.; Qian, T. Numerical study of the phase slip in two-dimensional superconducting strips. Physical Review B—Condensed Matter and Materials Physics 2008, 77 (17), 174517.

(106) Kulić, I.; Schiessel, H. Chromatin dynamics: Nucleosomes go mobile through twist defects. Physical review letters 2003, 91 (14), 148103.

(107) Richmond, T. J.; Davey, C. A. The structure of DNA in the nucleosome core. Nature 2003, 423 (6936), 145–150.

(108) Suto, R. K.; Edayathumangalam, R. S.; White, C. L.; Melander, C.; Gottesfeld, J. M.; Dervan, P. B.; Luger, K. Crystal structures of nucleosome core particles in complex with minor groove DNA-binding ligands. Journal of molecular biology 2003, 326 (2), 371–380.

(109) Gottesfeld, J. M.; Belitsky, J. M.; Melander, C.; Dervan, P. B.; Luger, K. Blocking transcription through a nucleosome with synthetic DNA ligands. Journal of molecular biology 2002, 321 (2), 249–263.

(110) Winger, J.; Nodelman, I. M.; Levendosky, R. F.; Bowman, G. D. A twist defect mechanism for atp-dependent translocation of nucleosomal DNA. Elife 2018, 7, e34100.

(111) Sabantsev, A.; Levendosky, R. F.; Zhuang, X.; Bowman, G. D.; Deindl, S. Direct observation of coordinated DNA movements on the nucleosome during chromatin remodelling. Nature communications 2019, 10 (1), 1720.

(112) Li, M.; Xia, X.; Tian, Y.; Jia, Q.; Liu, X.; Lu, Y.; Li, M.; Li, X.; Chen, Z. Mechanism of DNA translocation underlying chromatin remodelling by snf2. Nature 2019, 567 (7748), 409–413.

(113) Lorch, Y.; Davis, B.; Kornberg, R. D. Chromatin remodeling by DNA bending, not twisting. Proceedings of the National Academy of Sciences 2005, 102 (5), 1329–1332.

(114) Strohner, R.; Wachsmuth, M.; Dachauer, K.; Mazurkiewicz, J.; Hochstatter, J.; Rippe, K.; Längst, G. A’loop recapture’mechanism for acf-dependent nucleosome remodeling. Nature structural & molecular biology 2005, 12 (8), 683–690.

(115) Pande, V. S.; Beauchamp, K.; Bowman, G. R. Everything you wanted to know about markov state models but were afraid to ask. Methods 2010, 52 (1), 99–105.

(116) Husic, B. E.; Pande, V. S. Markov state models: From an art to a science. Journal of the American Chemical Society 2018, 140 (7), 2386–2396.

(117) Michael, A. K.; Grand, R. S.; Isbel, L.; Cavadini, S.; Kozicka, Z.; Kempf, G.; Bunker, R. D.; Schenk, A. D.; Graff-Meyer, A.; Pathare, G. R. Mechanisms of oct4-sox2 motif readout on nucleosomes. Science 2020, 368 (6498), 1460–1465.

(118) Liu, X.; Li, M.; Xia, X.; Li, X.; Chen, Z. Mechanism of chromatin remodelling revealed by the snf2-nucleosome structure. Nature 2017, 544 (7651), 440–445.

