## Supplementary Information for "Unveiling Nucleosome Dynamics: A Comparative Study Using All-Atom and Coarse-Grained Simulations Enhanced by Principal Component Analysis"

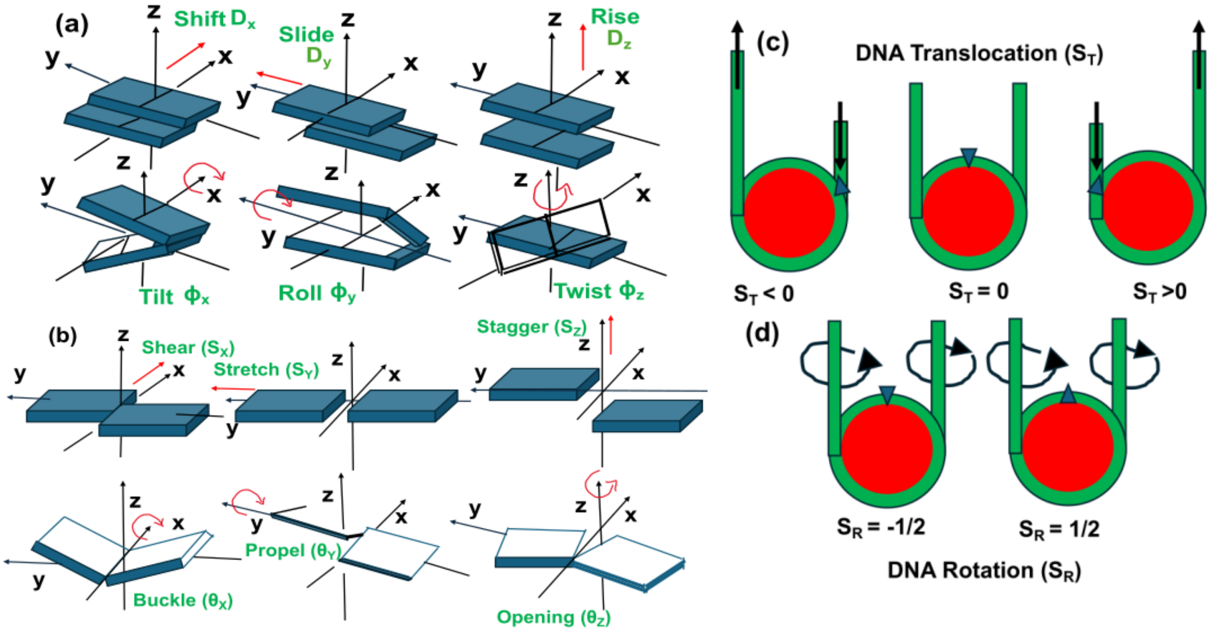


Figure S1 Schematic of (a) Inter-base pair parameters, (b) Intra-base pair parameters. Order parameter to characterize (c) translocation movement of nucleosome around histone, $S_T$ and (d) rotational movement of nucleosome relative to the histone, $S_R$ .

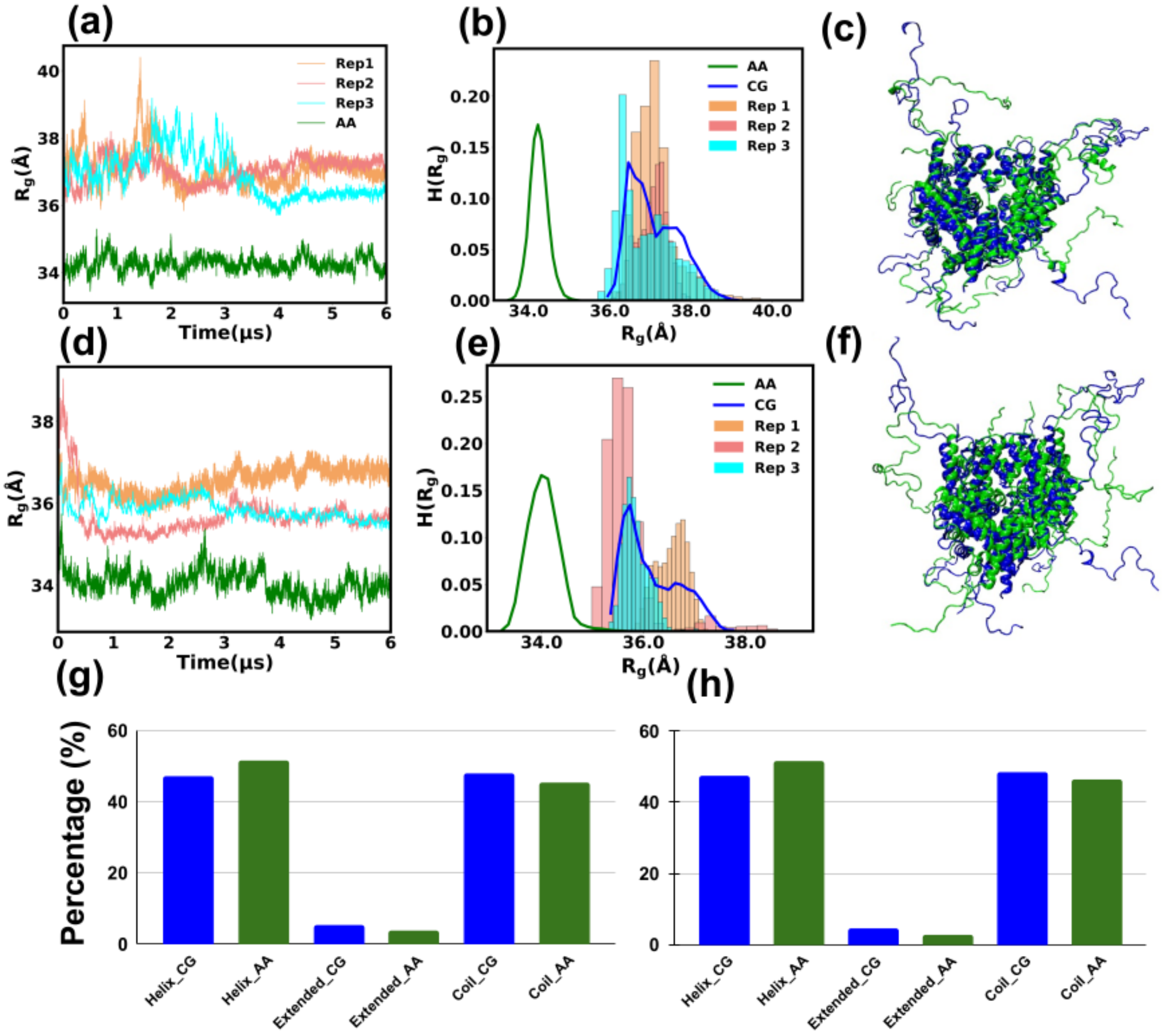


Figure S2: (a) Time evolution of radius of gyration ($R_g$) of protein considering C-alpha atom for ASP sequence. Results for three different replicas for coarse-grained trajectory and all atom trajectories are shown. (b) Histogram of $R_g$ for different replicas and atomistic data. (c) Representative structure for ASP histone sequence. (d) Time evolution of $R_g$ of histone for widom 601 sequence. Both CG replicas and atomistic simulation data is present. (e) Histogram of $R_g$ for widom 601 sequence. (f) Representative overlapped structure for widom 601 histone. Blue represent backmap atomic structure of CG trajectory while green represent structure obtained using atomistic simulations. Comparison of secondary structure propensity for (g) ASP sequence and (h) widom601 sequence.

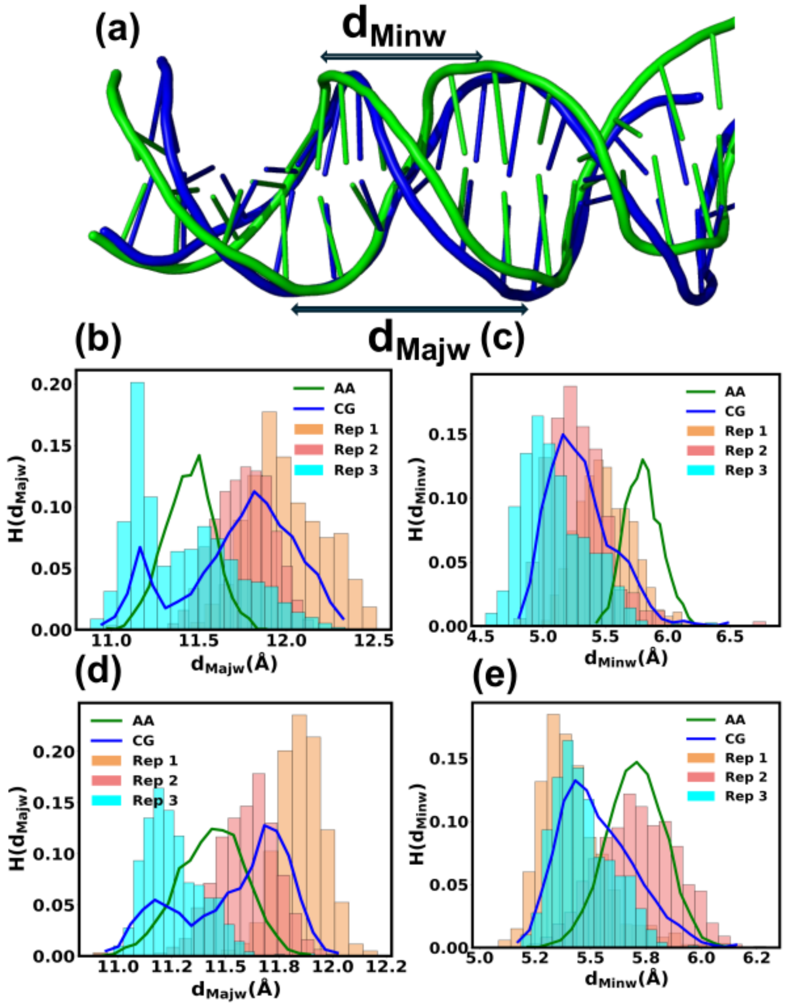


Figure S3 (a) Schematic of DNA major and minor width over the overlapped simulated structure. Blue represents backmap atomic structure of CG trajectory while green represent structure obtained using atomistic simulations. Histogram of DNA (b) major grove width and (c) minor groove width for ASP sequence. Results for both atomistic and CG replicas are shown in the figure. Histogram of DNA (d) major grove width and (e) minor groove width for widom 601 sequence.

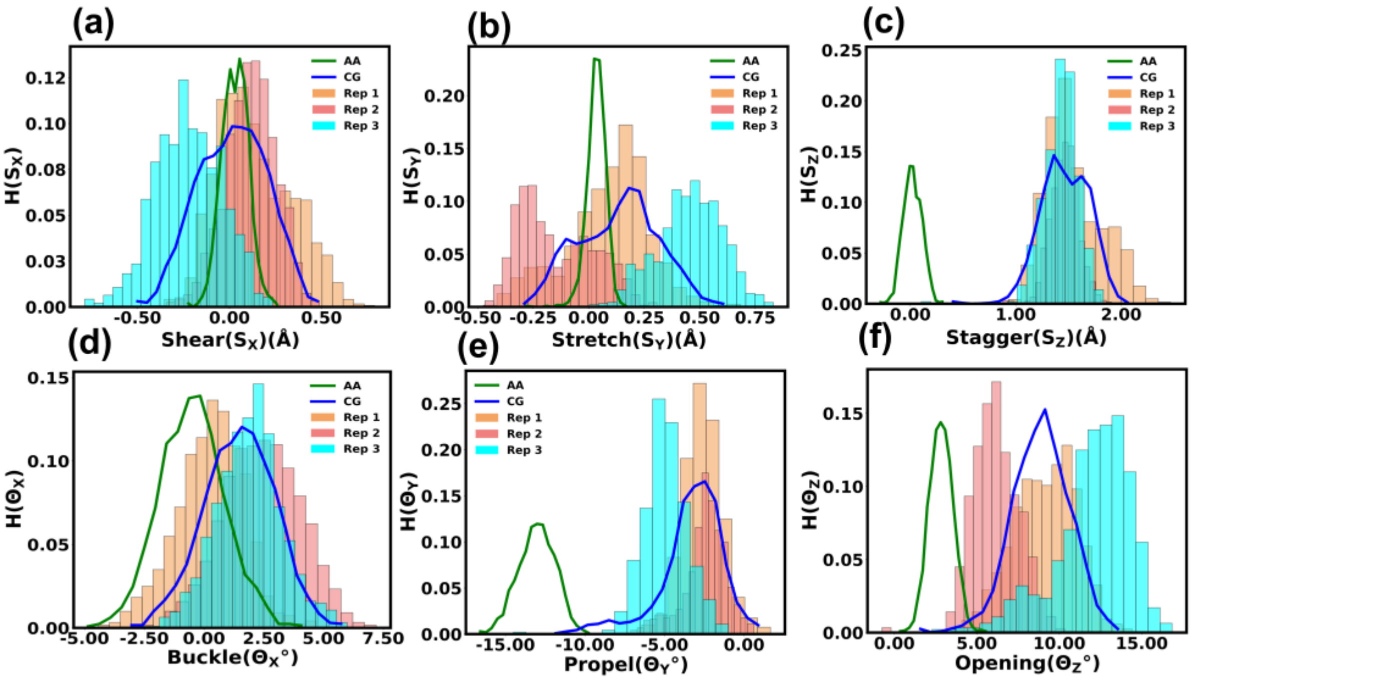


Figure S4 The histogram of DNA intra-base pair parameters for ASP sequence: (a) Shear, (b) Stretch, (c) Stagger, (d) Buckle, (e) Propel, (f) Opening. Results for both atomistic and three different CG replicas are shown in the figure.

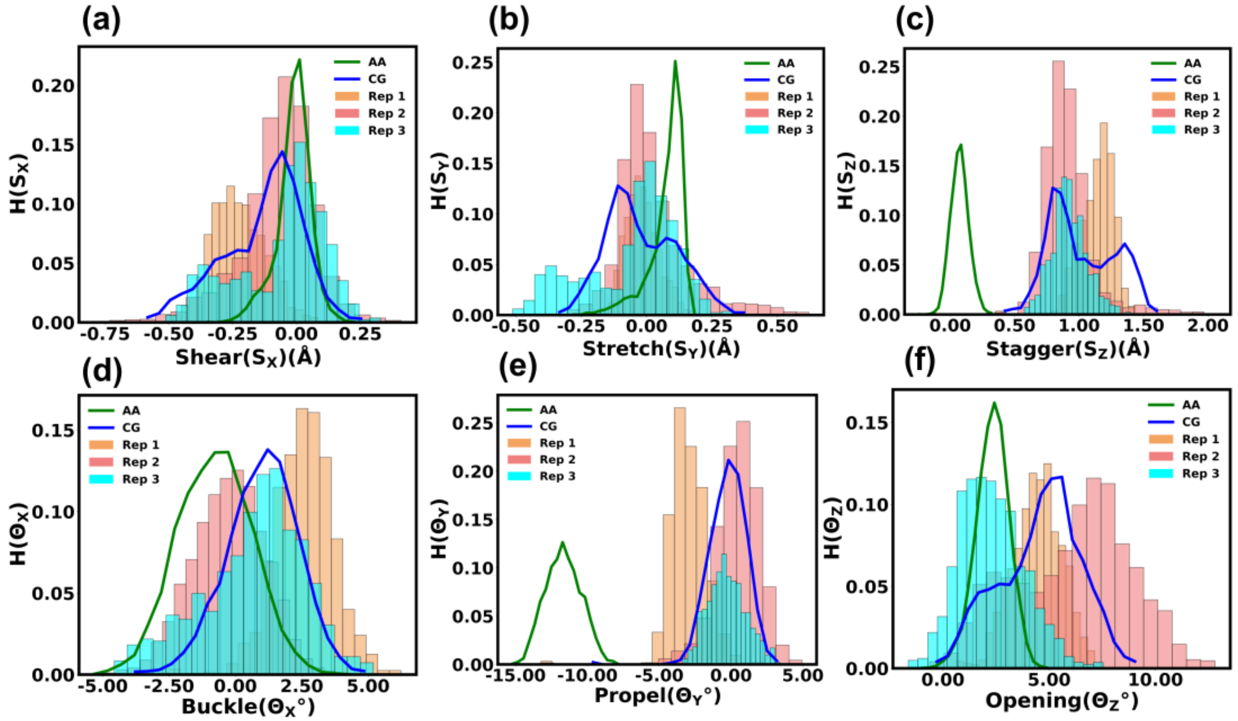


Figure S5 The histogram of DNA intra-base pair parameters for widom601 sequence: (a) Shear, (b) Stretch, (c) Stagger, (d) Buckle, (e) Propel, (f) Opening. Results for both atomistic and three different CG replicas are shown in the figure.

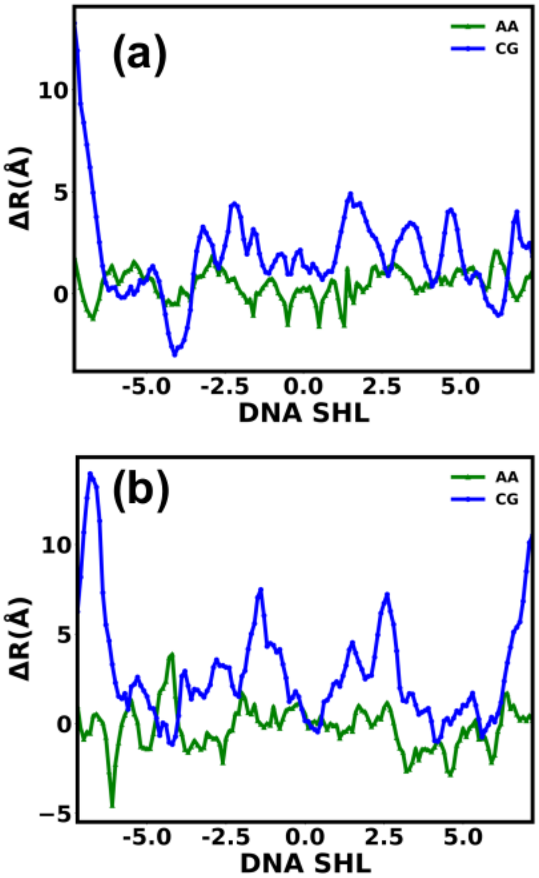


Figure S6 Change in average distance of each DNA base pair center represented in SHL notation over the simulated trajectory compared to crystal structure for (a) ASP and (b) Widom601 sequence.
